# Hippocampal protein aggregation signatures fully distinguish pathogenic and wildtype *UBQLN2* in amyotrophic lateral sclerosis

**DOI:** 10.1101/2022.01.12.475792

**Authors:** Kyrah M. Thumbadoo, Birger V. Dieriks, Helen C. Murray, Molly E. V. Swanson, Ji Hun Yoo, Nasim F. Mehrabi, Clinton Turner, Michael Dragunow, Richard L. M. Faull, Maurice A. Curtis, Teepu Siddique, Christopher E. Shaw, Lyndal Henden, Kelly L. Williams, Garth A. Nicholson, Emma L. Scotter

**Affiliations:** School of Biological Sciences, University of Auckland, Auckland, New Zealand; Centre for Brain Research, University of Auckland, Auckland, New Zealand; Department of Anatomy and Medical Imaging, University of Auckland, Auckland, New Zealand; Department of Pharmacology and Clinical Pharmacology, University of Auckland, Auckland, New Zealand; Department of Anatomical Pathology, LabPlus, Auckland City Hospital, Auckland, New Zealand; Departments of Neurology, Cell and Developmental Biology and Pathology, Northwestern University Feinberg School of Medicine, Chicago, USA; UK Dementia Research Institute Centre, Institute of Psychiatry, Psychology and Neuroscience, King’s College London, United Kingdom; Macquarie University Centre for Motor Neuron Disease Research, Macquarie Medical School, Faculty of Medicine, Health and Human Sciences, Macquarie University, Sydney, New South Wales, Australia; Northcott Neuroscience Laboratory, Australian and New Zealand Army Corps (ANZAC) Research Institute, Concord, New South Wales, Australia; Faculty of Medicine, University of Sydney, Sydney, New South Wales, Australia; Molecular Medicine Laboratory, Concord Repatriation General Hospital, Concord, New South Wales, Australia

**Keywords:** *UBQLN2*, ubiquilin 2, amyotrophic lateral sclerosis (ALS), frontotemporal dementia (FTD), neuropathology, hippocampus, human, TDP-43, polyGP, polyGA

## Abstract

Mutations in the *UBQLN2* gene cause X-linked dominant amyotrophic lateral sclerosis (ALS) and/or frontotemporal dementia (FTD) characterised by ubiquilin 2 aggregates in neurons of the motor cortex, hippocampus, and spinal cord. However, ubiquilin 2 neuropathology is also seen in sporadic and familial ALS or FTD cases not caused by *UBQLN2* mutations, particularly *C9orf72*-linked cases. This makes the mechanistic role of ubiquilin 2 mutations and the value of ubiquilin 2 pathology for predicting genotype unclear. Here we examine a cohort of 41 genotypically diverse ALS cases with or without FTD, including five cases with *UBQLN2* variants (resulting in p.S222G, p.P497H, p.P506S, and two cases with p.T487I). Using multiplexed (5-label) fluorescent immunohistochemistry, we mapped the co-localisation of ubiquilin 2 with phosphorylated TDP-43 (pTDP-43), dipeptide repeat aggregates, and p62, in the hippocampus of controls (n=5), or ALS with or without FTD in sporadic (n=20), unknown familial (n=3), *SOD1*-linked (n=1), *FUS*-linked (n=1), *C9orf72*-linked (n=5), and *UBQLN2*-linked (n=5) cases. We differentiate between i) ubiquilin 2 aggregation together with pTDP-43 or dipeptide repeat proteins, and ii) ubiquilin 2 self-aggregation promoted by *UBQLN2* gene mutations that cause ALS/FTD. Overall, we describe a hippocampal protein aggregation signature that fully distinguishes mutant from wildtype ubiquilin 2 in ALS with or without FTD, whereby mutant ubiquilin 2 is more prone than wildtype to aggregate independently of driving factors. This neuropathological signature can be used to assess the pathogenicity of *UBQLN2* gene variants and to understand the mechanisms of *UBQLN2*-linked disease.

## Introduction

*UBQLN2* gene mutations are a rare cause of amyotrophic lateral sclerosis (ALS) with or without dementia, and the only known causal mutations on the X-chromosome. *UBQLN2* [NM_013444] encodes the ubiquilin 2 protein, the best studied of five human ubiquilins, which contains a unique PXX domain comprising 12 proline-rich tandem repeats [1,2]. Like other ALS-and FTD-linked degradation proteins (sequestosome 1/p62 (hereafter p62), valosin-containing protein, optineurin, and TANK-binding kinase 1), a key role for ubiquilin 2 is to bind ubiquitinated, misfolded, and aggregated protein cargos, triaging them between the proteasome and autophagy intracellular degradation pathways [3–6]. New evidence suggests that ubiquilin 2 is intrinsically prone to self-assembly and undergoes reversible liquid-liquid phase separation, allowing it to bind ubiquitin-labelled proteins within phase-separated stress granules. It is postulated that upon ubiquitinated protein binding, ubiquilin 2 comes out of the liquid-droplet phase, bringing its cargo from the stress granule for delivery to the proteasome; ubiquilin 2 is therefore also implicated in stress granule disassembly [7–10].

These roles at the interface between RNA processing and protein degradation — two key pathways in ALS and FTD pathogenesis — may underpin why ubiquilin 2 mutations cause disease. ALS and/or FTD (ALS/FTD)-causing *UBQLN2* mutations that alter residues flanking and within the PXX domain impair ubiquilin 2 binding to the proteasome and promote self-oligomerisation and liquid-to-solid rather than liquid-liquid phase transition [9,11–14]. The mutation c.1490C>A, resulting in p.P497H, was the first disease-causing *UBQLN2* mutation identified, leading to the discovery in unrelated ALS/FTD families of mutations c.1489C>T (p.P497S), c.1516C>A (p.P506T), c.1525C>T (p.P509S) and c.1573C>T (p.P525S) all altering the PXX domain [1]. A c.1516C>T mutation, resulting in p.P506S also within the PXX domain, was later identified in a family with young onset ALS, ALS with FTD onset, or pure spastic paraplegia [15] while a c.1460C>T mutation, resulting in p.T487I just upstream of the PXX region, was found in two families with members in New Zealand and Australia [16,17]. *UBQLN2* mutations/variants have also been identified in ALS/FTD cases altering other residues within or flanking the PXX domain; in the ubiquitin-like domain; the stress-induced protein 1-like domains; or outside of known domains [18–37]. However, it is currently uncertain whether all of these *UBQLN2* variants are pathogenic (disease-causing).

The neuropathology of ALS/FTD caused by pathogenic *UBQLN2* mutations is characterised by aggregated ubiquilin 2 in the motor cortex, spinal cord, and hippocampus [1,15,17,23,38]. We previously reported ubiquilin 2-positive aggregates in the hippocampus of an individual with p.T487I *UBQLN2*-linked ALS+FTD, but not in sporadic ALS [38]. However, ubiquilin 2 inclusions are not specific to *UBQLN2*-linked ALS/FTD cases and have been identified previously in sporadic and familial ALS, and ALS-dementia, regardless of whether ubiquilin 2 is wildtype or mutant [1,15,17,23,38,39]. Some of these ubiquilin 2 inclusions are immunopositive for other ALS/FTD-linked proteins such as TDP-43 or phosphorylated TDP-43 (pTDP-43), FUS, p62, optineurin, or ubiquitin [1,17,40–44]. In addition, deposition of ubiquilin 2 in the hippocampus is a characteristic feature of ALS/FTD caused by *C9orf72* hexanucleotide repeat expansions, in which ubiquilin 2 is seen in the presence of dipeptide repeat (DPR) proteins [39]. Thus, while ubiquilin 2 is clearly involved in ALS/FTD pathogenesis regardless of aetiology, the role of *UBQLN2* mutations and the predictive value of ubiquilin 2 labelling in relation to genotype is still unclear.

There is a clear need to determine whether *UBQLN2* mutations cause ubiquilin 2 neuropathology that is distinct from the wildtype ubiquilin 2 neuropathology that is seen in ALS/FTD with other genotypes. If they do, then defining a pathological signature for *UBQLN2*-linked disease will provide mechanistic insights and aid assessment of the pathogenicity of *UBQLN2* genetic variants which are not yet confirmed to be causative. Because ubiquilin 2 aggregation across a range of ALS/FTD genotypes occurs in the hippocampus, this region may provide insight into the requirements for mutant and wildtype ubiquilin 2 aggregation. Here we map the hippocampal ubiquilin 2 protein deposition signature with respect to pTDP-43, the two most abundant *C9orf72-*linked DPR proteins, and p62, in ALS/FTD with and without *UBQLN2* mutation.

## Materials and methods

### Systematic review of *UBQLN2-*linked ALS/FTD neuropathology

Journal articles were identified using PubMed that were published between Jan 1993 and May 2023. Search terms were *UBQLN2*, ubiquilin 2, amyotrophic lateral sclerosis, and ALS, combined with neuropathology, tissue, or immunohistochemistry. Articles which did not contain ubiquilin 2 neuropathological information in post-mortem ALS/FTD human brain tissue, were not primary research articles, or were not published in English were excluded. Seven papers were identified from which neuropathological data for TDP-43, ubiquilin 2, ubiquitin, p62, *C9orf72*-linked DPR proteins, FUS, and SOD1 protein aggregates in the spinal cord and hippocampus were extracted and tabulated.

### Patient demographics and hippocampal brain tissue

Formalin-fixed paraffin-embedded (FFPE) post-mortem hippocampal tissue from 6 neurologically normal and 30 ALS cases with or without FTD, processed as previously described [45] were obtained from the Neurological Foundation Human Brain Bank at the Centre for Brain Research, Auckland, New Zealand. These included twenty cases with sporadic ALS (one with co-morbid FTD), three with familial ALS of unknown genotype, one with *SOD1*-linked ALS (p.E101G), five with *C9orf72*-linked ALS, and one with ALS+FTD with the *UBQLN2* p.T487I mutation (pedigree ID FALS5 IV:18 in Fig. 1A of [17] and in this report; coded MN17 in Figure 1D of [38] and in this report), giving a total cohort of 41 cases. Note that MN17 was stored in fixative for 7 years before embedding. All non-*SOD1*-linked NZ cases had confirmed pTDP-43 proteinopathy in the motor cortex. FFPE hippocampal tissue was also obtained from: the Victoria Brain Bank from a relative of the *UBQLN2* p.T487I case (p.T487I, ALS+FTD; pedigree ID V:7 in Fig. 1A of [17] and in this report); the London Neurodegenerative Diseases Brain Bank and Brains for Dementia from an unrelated *UBQLN2*-linked case (p.P506S, ALS+FTD:, pedigree ID II.2 in Fig. 1B of [15]), and a case with a *UBQLN2* variant of unknown significance (p.S222G, progressive supranuclear palsy; Supplemental_variant_data of [46]), and a *FUS*-linked case (p.P525L, ALS); and from the Northwestern University Feinberg School of Medicine, USA from another unrelated *UBQLN2*-linked case (p.P497H, ALS; pedigree ID V:I of Family #186 in Fig. 1A of [1]). All clinical and neuropathological diagnoses were conducted as described previously [1,15,17,38]. Patient demographic and clinical information is summarised in Supplementary Table 1.

**Figure 1.**
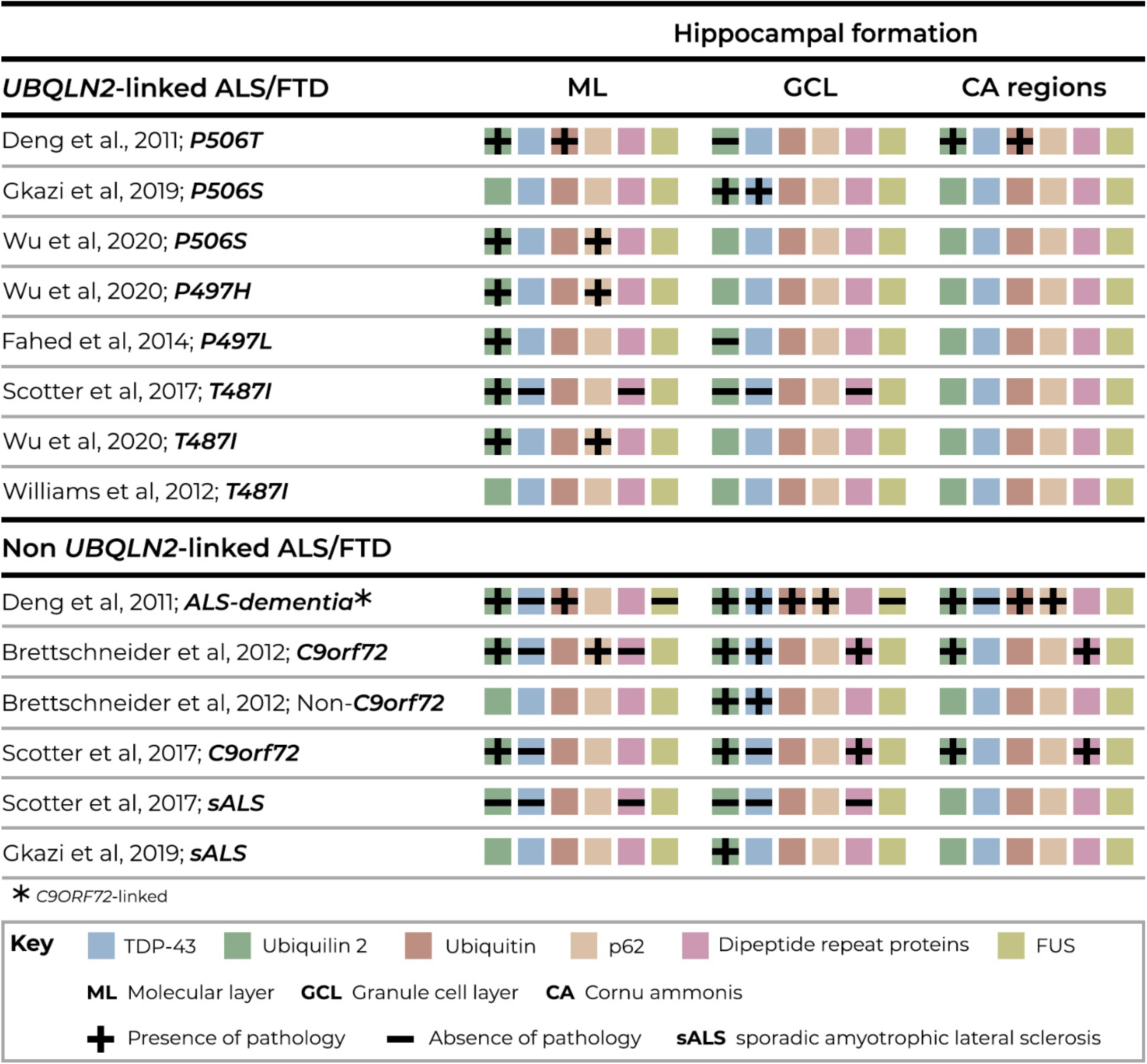
Ubiquilin 2 labelling in previous studies fails to discriminate between *UBQLN2*-linked and other genotypes of ALS/FTD. Previously published immunohistochemical analyses of hippocampus molecular layer (ML), granule cell layer (GCL), and cornus ammonus (CA) regions fail to discriminate between *UBQLN2-*linked ALS/FTD and ALS/FTD with other genotypes. Key shown within figure. Hippocampal ubiquilin 2 (green) pathology is reported when ubiquilin 2 is either mutant or wildtype; being present in the molecular layer in *UBQLN2*-linked and *C9orf72*-linked ALS/FTD (including “ALS-dementia”) but not in sALS; and in the granule cell layer in *C9orf72*-linked ALS/FTD (including “ALS-dementia”). Hippocampal molecular layer ubiquilin 2 (green) is frequently reported and found together with ubiquitin (orange) or p62 (beige), but not TDP-43 (blue). Hippocampal granule cell layer ubiquilin 2 pathology is found together with dipeptide repeats (DPRs, pink) with or without TDP-43; but is variably present in *UBQLN2*-linked ALS/FTD and sALS. In the hippocampal CA regions, ubiquilin 2 is present in *UBQLN2*-linked and in *C9orf72*-linked ALS/FTD (including “ALS-dementia”) where it is found together with DPRs, ubiquitin and p62.

### Fluorescent immunohistochemistry and image acquisition

#### Multiplex fluorescent immunohistochemistry

Multiplex (five-label) immunohistochemistry was performed as described previously [47–49]. Briefly, tissue sections were cut with a microtome in the transverse plane at a thickness of 7-10 µm and mounted onto Superfrost Plus slides (Thermo Fisher Scientific). Mounted sections were dried at room temperature for a minimum of 1 week before immunohistochemistry. Slides were heated to 60 °C for 1 h on a hot plate, then dewaxed and rehydrated through a xylene-alcohol-water series: 100% xylene, 2x 30 min; 100% ethanol, 2x 15 min; 95%, 85%, 75% ethanol, 5 min each; water 3x 5 min. Antigens were retrieved through immersion in 10 mM sodium citrate buffer (0.05% Tween 20, pH 6.0) in a pressure cooker (Retriever 2100, Electron Microscopy Sciences) at 120 °C for 20 min and cooled to room temperature for 100 min. Sections were washed 3x 5 min in 1x phosphate-buffered saline (PBS) and wax borders were drawn with an ImmEdge Hydrophobic Barrier PAP pen. Sections were permeabilised in PBS-T (PBS with 0.2% Triton™ X100) for 15 min at 4 °C, followed by 3x 5-min PBS washes. Lipofuscin autofluorescence was quenched using TrueBlack® Lipofuscin quencher (Biotium) 1:20 in 70% ethanol for at least 30 seconds followed by three vigorous water washes. Sections were blocked with 10% normal donkey serum (Thermo Fisher Scientific) in PBS for 1 h before 4 °C overnight incubation with primary antibodies targeting ubiquilin 2, phosphorylated TDP-43 (pTDP-43), p62, and *C9orf72-*linked dipeptide repeat proteins polyGA and polyGP (Supplementary Table 2). Following PBS washes, species-and isotype-specific secondary antibodies and Hoechst 33342 nuclear stain (Supplementary Table 3) were applied for 3 h in 1% normal donkey serum at room temperature. After final 3x 5-min PBS washes, sections were coverslipped with #1.5 coverslips (Menzel-Gläser) using ProLong™ Gold Antifade Mountant (Thermo Fisher Scientific). All primary antibodies used in this study have been previously validated in human brain tissue [6,38,50,51], and secondary antibodies showed no non-specific binding or signal bleed-through into adjacent channels (Supplementary Fig. 1) or cross-reactivity (Supplementary Fig. 2).

Multiplex (five-label) images were acquired using a Zeiss Z2 Axioimager with a MetaSystems VSlide slide scanning microscope (20x dry magnification lens, 0.9 NA) with a Colibri 7 solid-state fluorescent light source. Sections were imaged with the same exposure time and gain settings for each staining combination where possible. *UBQLN2*-linked ALS/FTD case V:7-p.T487I showed consistently poor Hoechst immunoreactivity, while *UBQLN2*-linked ALS/FTD case MN17-p.T487I showed poor immunoreactivity overall, likely due to long-term fixation, therefore a longer exposure was used for both cases when imaging. Single filters, as described and validated previously [49], were used to excite fluorophores and detect the following wavelengths: Filter set #1 (LED 385; Em 447/60 nm), #3 (LED 475; Em 527/20 nm), #4 (LED 555; Em 580/23 nm), #5 (LED 590; Em 628/32 nm), #6 (LED 630; Em 676/29 nm), and #7 (LED 735; Em 809/81 nm) and visualised with MetaFer software (MetaSystems, v.3.12.1) equipped with a CoolCube 4m TEC (monochrome) sCMOS digital camera. Image tiles were seamlessly stitched using MetaCyte software, and stitched images were extracted using VSViewer software (MetaSystems, v.1.1.106).

#### Double-label immunohistochemistry

For super-resolution stimulated emission depleted (STED) microscopy of *C9orf72*-linked ALS case MN28, double-label immunohistochemistry was performed. Immunohistochemistry was performed as above with primary antibody targeting ubiquilin 2 and detected by goat anti-mouse IgG_2a_ Alexa Fluor 594, but with primary antibody targeting polyGA detected by goat anti-mouse IgG_1_ Biotin (A10519, Thermo Fisher Scientific), followed by 3x 5-min PBS washes, and an additional incubation with Star Red neutravidin (STRED-0121, Abberior) for 1 hour at room temperature. After final 3x 5-min PBS washes, sections were coverslipped with #1.5 coverslips using ProLong™ Glass Antifade Mountant (P36984, Thermo Fisher Scientific).

STED images were acquired using an Abberior Facility STED microscope (60x UPLXAPO oil immersion lens, 1.42 NA) using ImSpector Lightbox software (Specim, v.16.3.13779). A 561-nm pulsed diode laser was used to excite Alexa Fluor 594, and a 640-nm diode laser was used to excite Alexa Fluor 647. For STED imaging, a pulsed 775-nm laser was used for depletion of both fluorophores. After scanning, the images were processed using the PureDenoise plugin [52] for ImageJ (National Institutes of Health, USA v1.53f51).

Final figures of multiplex and double-label immunohistochemistry were compiled using Adobe Photoshop CC (Adobe Systems Incorporated, v24.4.1). For clarity, multiplex labels are shown in separate panels, either pTDP-43, ubiquilin 2, and p62, or ubiquilin 2, polyGA, and polyGP, with negative labelling not shown.

#### Relatedness analysis

To further examine the p.T487I mutation in ALS/FTD, genome-wide genotype data from the family of *UBQLN2*-linked ALS+FTD case MN17 (FALS5) and another Australian family with an identical p.T487I mutation (FALS14) were analysed to determine whether they inherited the mutation from a common ancestor (pedigree in Supplementary Fig. 3, sample information in Supplementary Table 4). These two families were previously reported to share a haplotype identical-by-state over the *UBQLN2* locus [17], but genealogy analysis had been unable to link the pedigrees. Three individuals from FALS5 (all affected, pedigree IDs III:8, IV:9, IV:18 (MN17)) and two individuals from FALS14 (pedigree IDs II:1 (unaffected) and III:2 (affected)) underwent SNP genotyping using the Illumina Infinium Global Screening Array v2.0. Identity-by-descent (IBD) analysis was performed using XIBD software [53] with the combined HapMap Phase II and III European (CEU) cohort as a reference dataset. SNPs in high linkage disequilibrium (r^2^> 0.95) or SNPs with a low minor allele frequency (MAF < 0.01) were removed from analysis, as well as SNPs with missing genotype calls in two or more samples. 39,002 SNPs remained for analysis and 211 IBD segments greater than 3 cM were identified. The degree of relationship was estimated for each pair of samples using the lengths of inferred IBD segments as in Estimation of Recent Shared Ancestry [54].

## Results

### Ubiquilin 2 labelling in previous studies failed to discriminate between *UBQLN2-***linked and other genotypes of ALS/FTD**

Systematic review of the literature describing ubiquilin 2 neuropathology in human ALS/FTD identified 137 results, of which 7 articles met the criteria for full review. The hippocampal neuropathology of ubiquilin 2 with respect to five other ALS/FTD-linked proteins is summarised in Fig. 1. Ubiquilin 2 aggregate deposition in the hippocampal molecular layer was most frequently reported upon, being found in *UBQLN2*-linked and *C9orf72*-linked ALS/FTD, sometimes together with ubiquitin or p62 [6]. Ubiquilin 2 aggregate deposition in the granule cell layer of the hippocampal dentate gyrus (GCL) was inconsistent between cases of *UBQLN2*-linked ALS/FTD but was found in all cases of “ALS-dementia” in [1] (all later confirmed to be *C9orf72*-positive) and in a single sporadic ALS/FTD case [15]. Cornu ammonis (CA) ubiquilin 2 pathology had been reported in both *UBQLN2*-linked and *C9orf72*-linked ALS/FTD [1,38,39]. Overall, neither ubiquilin 2 staining alone nor in combinations previously tested could distinguish mutant ubiquilin 2 aggregation in *UBQLN2*-linked ALS/FTD from wildtype ubiquilin 2 aggregation in other ALS/FTD genotypes.

### Ubiquilin 2 and other pathology in ALS/FTD of various genotypes

#### Control and familial ALS cases had no hippocampal ubiquilin 2 aggregates but certain cases had sparse p62-only aggregates

Five of the six non-neurodegenerative disease control cases included in this study showed no deposition of ubiquilin 2, pTDP-43, p62, or DPR protein aggregates in any region of the hippocampus (not shown). The remaining control case (H230) showed sparse p62 aggregates in the GCL and CA pyramidal cell layer (CA – pyr) (Fig. 1A_1,2_).

All five familial ALS cases also showed sparse p62 aggregates in the GCL and CA – pyr regions; three cases were of unknown genotype, represented by case MN14 (Fig. 1B_1,2_), one harboured a *SOD1* p.E101G mutation (Fig. 1C_1,2_), and one harboured a *FUS* p.P525L mutation (Fig. 1D_1,2_). No other pathology was observed in familial cases in hippocampal regions.

#### Sporadic ALS cases had no hippocampal ubiquilin 2 aggregates but certain cases had pTDP-43-p62 aggregates

None of the 20 sporadic ALS cases (one with co-morbid FTD) showed hippocampal ubiquilin 2 or DPR aggregates (not shown). Fourteen cases (70%) were devoid of either ubiquilin 2 or pTDP-43 in the GCL, ML, CA-pyr, and CA – lm/rad layers (not shown), however 4 cases (20%) had compact p62-positive NCIs in the GCL and CA – pyr regions that were negative for all other markers (Supplementary Fig. 4). Six cases (30%; including representative cases MN15 with ALS+FTD, and MN30 with ALS) had compact or net-like perinuclear pTDP-43-positive neuronal cytoplasmic inclusions (NCIs) in the GCL and CA – pyr regions ranging in density from sparse to frequent, suggestive of stage 4 pTDP-43 proteinopathy [55]. These pTDP-43 aggregates were almost always p62-positive and ubiquilin 2-negative (Fig. 2E_1,2_ and F_1,2_, white arrows), with the exception of 1-2 perinuclear aggregates that were not p62-labelled (Fig. 2F_1_, yellow arrow).

**Figure 2.**
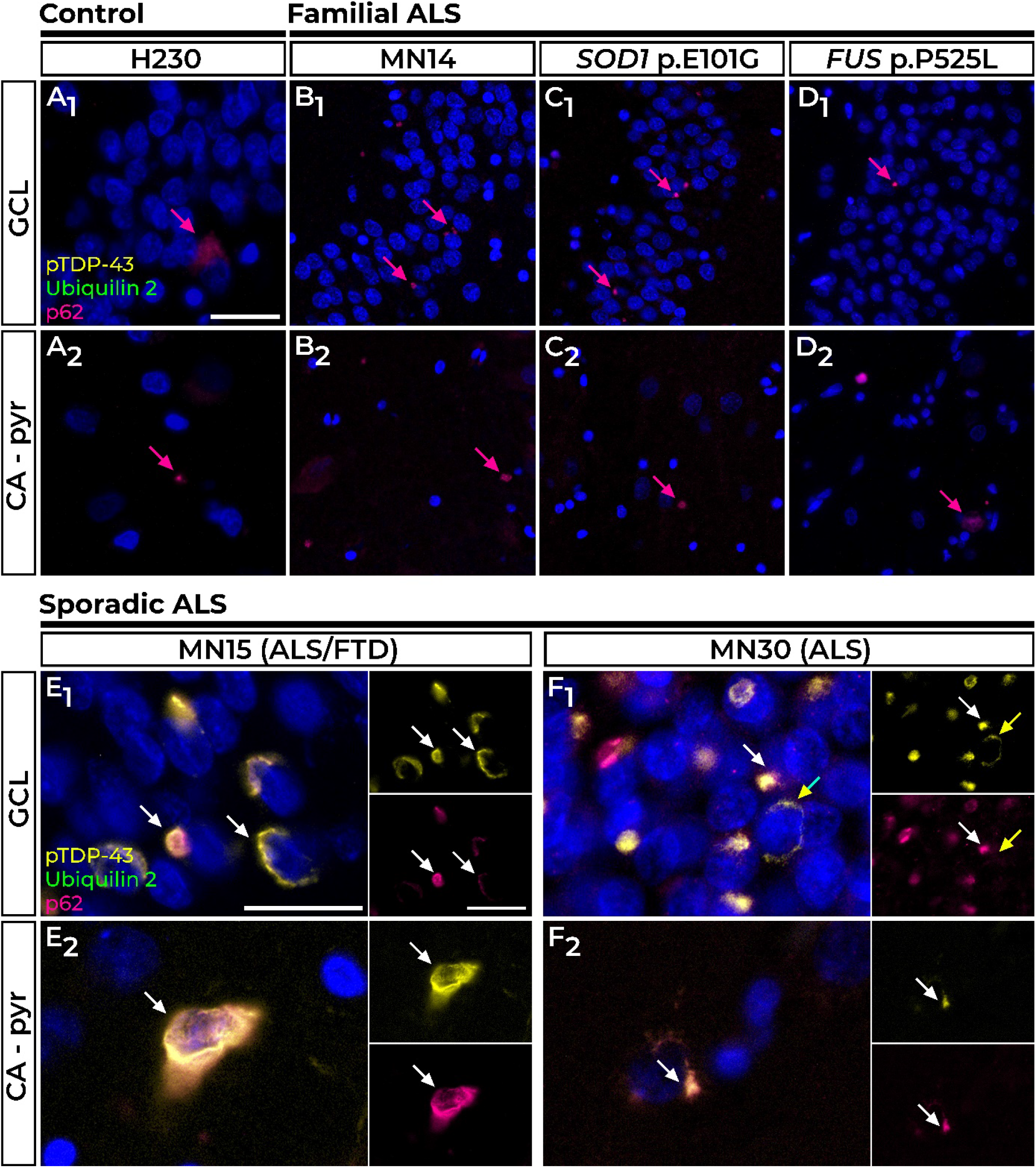
Pathology in the hippocampal dentate gyrus and cornu ammonis regions of control, familial, and sporadic cases. The majority of control tissue had no pTDP-43 or ubiquilin 2 pathology, but one case had sparse p62 immunoreactivity in the GCL and CA-pyr layers (**A** H230, pink arrows). Such punctate p62-positive inclusions were also found in the GCL (**B_1_-D_1_**), and CA-pyr layers (**B_2_-D_2_**) in MN14, an ALS case of unknown genotypic cause (**B**), MN24, a *SOD1* p.E101G case (**C**), and P525L, a *FUS* p.P525L case (**D**). P62-co-labelled perinuclear pTDP-43 aggregates negative for ubiquilin 2 were found in the GCL and CA-pyr regions of sporadic case ALS/FTD case MN15 (**E_1_** and **E_2_**, white arrows), and sporadic ALS case MN30 (**F_1_** and **F_2_**, white arrows). Rare pTDP-43 aggregates negative for p62 are indicated by the yellow arrow in **F_2_**. Scale bar A-D, 50 µm; E-F, 25 µm. Abbreviations: CA – pyr, cornu ammonis – pyramidal cells; GCL, granule cell layer.

#### C9orf72 cases had molecular layer ubiquilin 2-only aggregates, granule cell layer ubiquilin 2-DPR-p62 aggregates, and one case had granule cell layer ubiquilin 2-DPR-pTDP-43-p62 aggregates

Four of the five *C9orf72-*linked ALS cases showed identical hippocampal pathology (represented by case MN18), while the fifth case (MN28) differed only by having pTDP-43 aggregates in the GCL and CA – pyr, described below.

All five cases shared a signature of numerous ubiquilin 2 aggregates in the ML, GCL and CA – l-m/rad regions. In the ML and CA – l-m/rad, ubiquilin 2 aggregates were punctate or skein-like, dendritic, and very rarely co-localised with p62 (Fig. 3A_1,2_ and B_1,2_, green arrowheads) and never with DPR proteins (Fig. 3C_1,2_ and D_1,2_, green arrowheads). In the CA – pyr, aggregates did not contain ubiquilin 2 (Fig. 3A_3_-D_3_) but were positive for p62 (Fig. 3A_3_, B_3_, pink arrows), polyGA, and polyGP (Fig 3C_3_, D_3_, orange arrows and insets), with pTDP-43 in case MN28 also seen rarely (Fig. 3B_3_, yellow arrow).

**Figure 3.**
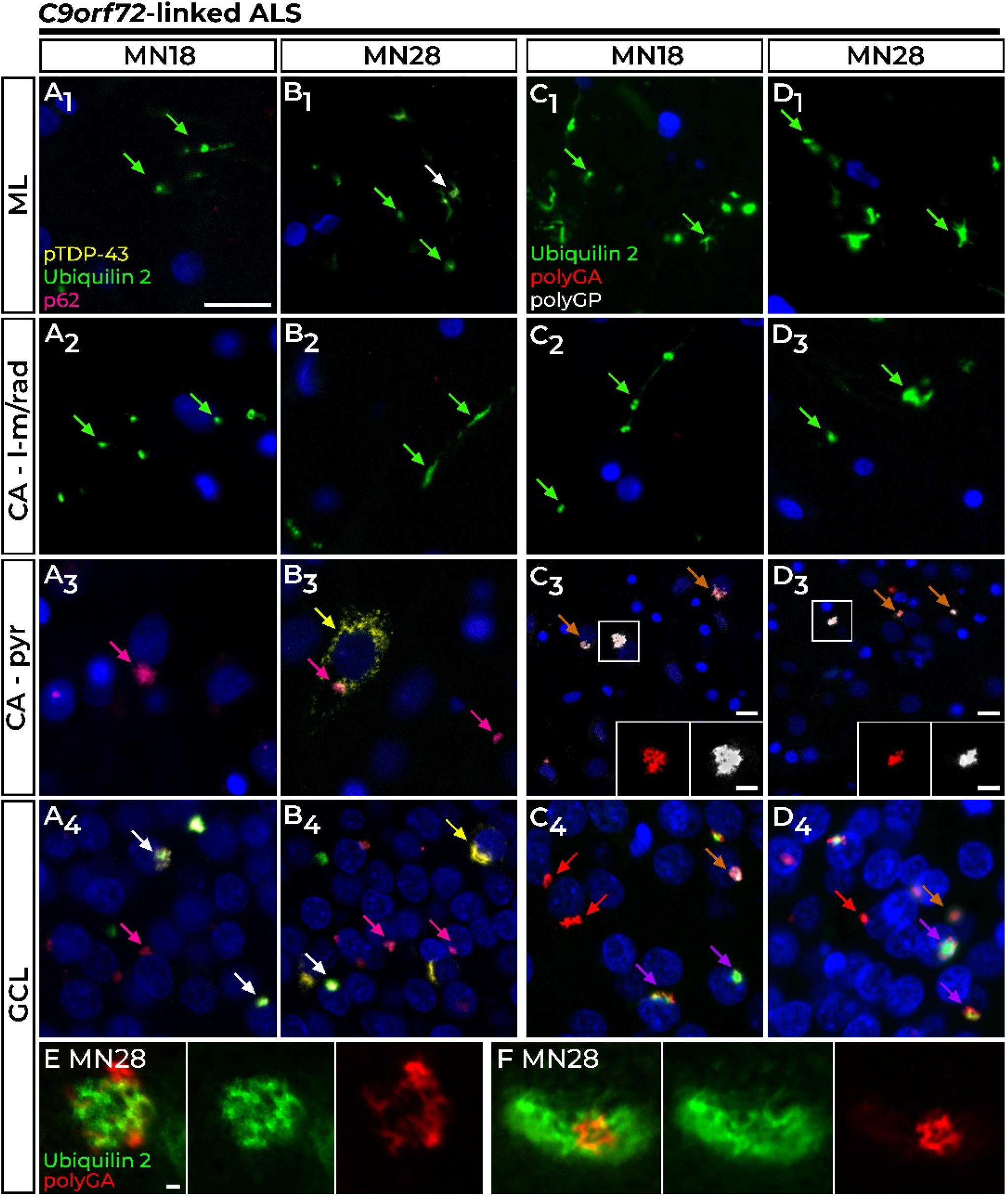
Pathology in the hippocampal dentate gyrus and cornu ammonis regions of *C9orf72*-linked cases. Wispy ubiquilin 2 aggregates in the ML and CA – l-m-rad layers were predominantly p62 negative (**A,B_1_** and **A,B_2_**, green arrows), with larger aggregates also co-labelling with p62 in the ML (**B_1_**, white arrow). In MN18, the CA – pyr cells harboured stellate p62 aggregates (**A_3_**, pink arrow) while the GCL showed stellate p62 aggregates either with ubiquilin 2 (**A_4_**, white arrows) or without (**A_4_**, pink arrow). Case MN28 shared this pattern of p62 and ubiquilin 2 pathology with the additional presence of perinuclear pTDP-43 inclusions, often found without other co-labelling in the CA-pyr region (**B_3_**, yellow arrow) and GCL (**B_4_**, yellow arrow). Ubiquilin 2 aggregates in the ML and CA – l-m/rad layers were negative for dipeptide repeat proteins (**C_1_-C_2_**, **D_1_-D_2_**, green arrows). The dominant pathology in CA – pyr regions was the presence of polyGA and polyGP inclusions negative for ubiquilin 2 (**C_3_** and **D_3_**, orange arrowheads). All GCL ubiquilin 2 was positive for at least one dipeptide repeat protein (**C_4_** and **D_4_**, purple arrows). Ubiquilin 2-negative aggregates were comprised of both dipeptide repeat proteins (**C_4_** and **D_4_**, orange arrows) or most commonly, polyGA without other co-labelling (**C_4_** and **D_4_**, red arrows). Super-resolution STED microscopy of co-localised ubiquilin 2-polyGA GCL aggregates in case MN28 demonstrated polyGA enmeshed with and encircling ubiquilin 2 (**E**) or ubiquilin 2 labelling around a core of polyGA (**F**). Scale bar in main images A-D, 50 µm; zooms in C_4_ and D_4_, 25 µm; in E-F, 1 µm. Abbreviations: CA – lm/rad, cornu ammonis – lacunosum-molecular and radiatum layers; CA – pyr, cornu ammonis – pyramidal cells; GCL, granule cell layer.

In the GCL, ubiquilin 2 aggregates were perinuclear NCIs that were negative for pTDP-43 except rare aggregates in case MN28 (Fig. 3B_4_, yellow arrow), but always positive for p62 (Fig. 3A_1_, B_1_, white arrows and yellow arrow) and poly(glycine-arginine) (polyGA), but rarely polyGP (Fig 3C_1_, D_1_, yellow arrows). Ubiquilin 2 aggregates positive for pTDP-43 were rare (1-2 GCL cells per section of MN28) and comprised a pTDP-43 ‘shell’ surrounding polyGA either with polyGP (Supplementary Fig. 5A, white arrow) or without polyGP (Supplementary Fig. 5A, purple arrow). This supports a previous report that ubiquilin 2 in *C9orf72*-linked cases rarely co-localises with GCL pTDP-43, doing so only when DPR proteins are present in the aggregate [56]. Therefore, wildtype ubiquilin 2 aggregation may be ‘seeded’ in the GCL by polyGA rather than by pTDP-43.

#### UBQLN2 cases show molecular layer ubiquilin 2-p62 aggregates and certain cases had granule cell layer pTDP-43 aggregates

All four *UBQLN2-*linked ALS/FTD cases showed signature ubiquilin 2-positive but pTDP-43-negative punctate aggregates in the hippocampal ML and CA – lm/rad regions, albeit at variable loads (Fig. 4A_2,3_, B_2,3_, C_2,3_, D_2,3_, white arrows). In the ML, ubiquilin 2 aggregates were punctate and appeared to be localised to the dendritic spines of the granule cells, as reported in *UBQLN2-*linked ALS/FTD and mutant *UBQLN2* rodent models [1,57]. In contrast to *C9orf72*-linked cases, these ML ubiquilin 2 aggregates almost always co-localised with p62. The PSP case with a *UBQLN2* p.S222G variant of unknown significance did not share this signature of ubiquilin 2-and p62-positive aggregates in the ML and CA – lm/rad, instead having p62-only aggregates in the CA – lm/rad (Fig. 4E_2,3_, pink arrowheads).

**Figure 4.**
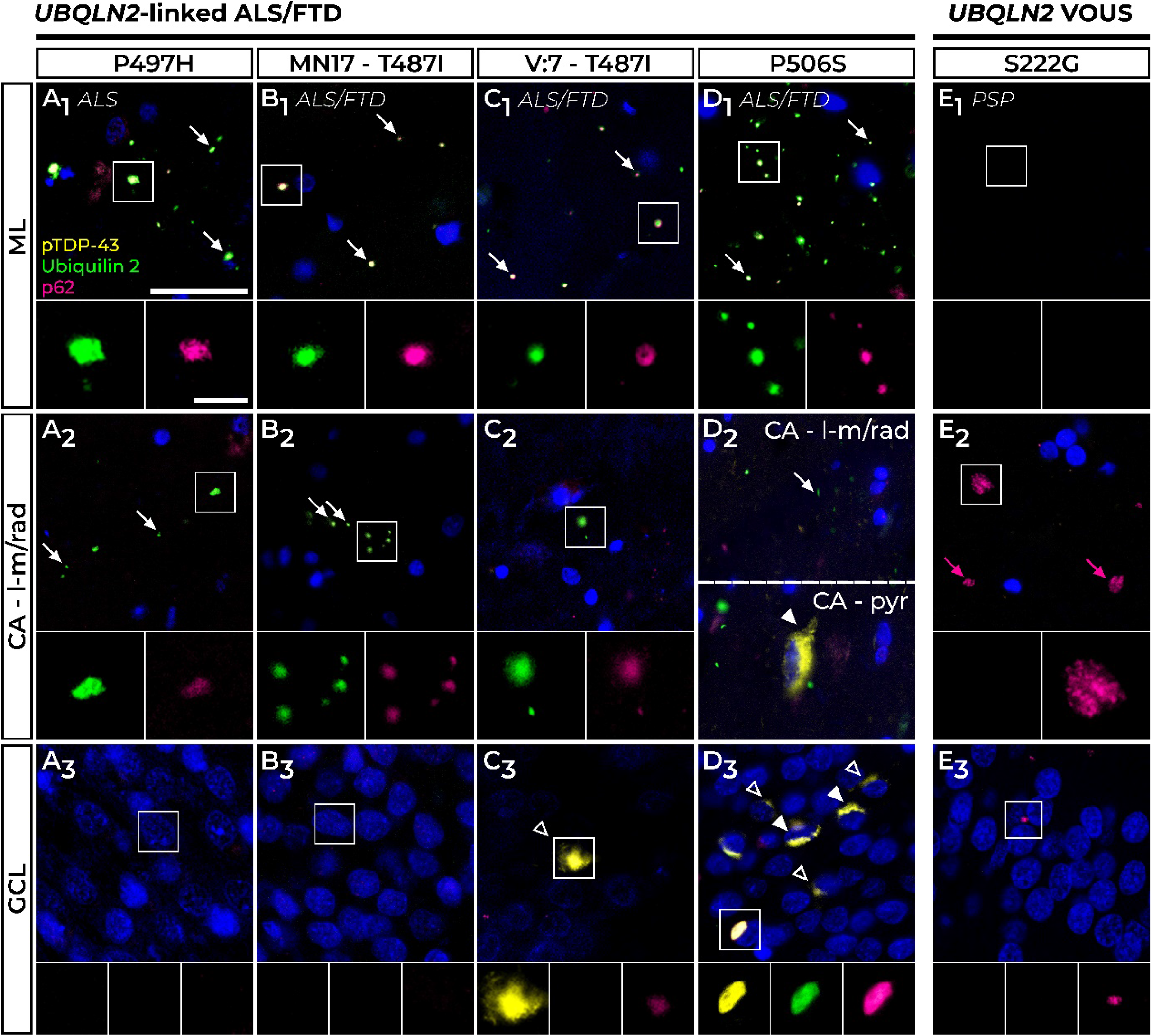
Ubiquilin 2, pTDP-43, and p62 pathology in the hippocampal dentate gyrus and cornu ammonis regions of *UBQLN2*-linked cases. All *UBQLN2*-linked ALS/FTD cases showed numerous p62-positive ubiquilin 2 aggregates, indicated by white arrows, in the ML (**A_1_**-**D_1_**) and CA – l-m/rad layers (**A_2_**-**D_2_**). Case P506S had the additional presence of aggregates positive for pTDP-43, ubiquilin 2, and p62 in the CA – pyr layer (**D_2_**, filled white arrowhead). Cases P497H and MN17-T487I showed no aggregate labelling in the GCL (**A_3_**, **B_3_**). ALS/FTD cases V:7 – T487I and P506S showed rare GCL p62-positive pTDP-43 aggregates negative for ubiquilin 2 (**C_3_**, **D_3_**, hollow white arrowheads). Only in P506S were the numerous compact pTDP-43 aggregates found in the GCL co-localised with ubiquilin 2 and p62 (**D_3_**, filled white arrowheads). A PSP case harbouring a *UBQLN2* p.S222G variant of uncertain significance (VOUS) (**E**) had pathology in the ML (**E_1_**), and sparse p62 positivity (pink arrows) in the CA – lm/rad layers (**E_2_**) and GCL (**E_3_**). Scale bar in main images, 25 µm; zooms, 5 µm. Abbreviations: CA – lm/rad, cornu ammonis – lacunosum-molecular and radiatum layers; CA – pyr, cornu ammonis – pyramidal cells; GCL, granule cell layer; PSP, progressive supranuclear palsy.

Confocal microscopy of the ML and CA – lm/rad regions confirmed that the p62-positive mutant ubiquilin 2 aggregates in *UBQLN2-*linked ALS/FTD cases (Supplementary Fig. 5A, representative p.P506S case shown) differed from wildtype ubiquilin 2 aggregates in those regions in *C9orf72-*linked ALS, which were predominantly p62 negative (Supplementary Fig. 5B, representative case MN28). Further, while mutant ubiquilin 2 in *UBQLN2*-linked cases formed small compact aggregates (Supplementary Fig. 6A) wildtype ubiquilin 2 in *C9orf72-*linked cases formed small compact aggregates and wispy skein-like structures (Supplementary Fig. 6B). Therefore, mutant but not wildtype ubiquilin 2 in the hippocampal ML and CA – lm/rad regions may promote or scaffold the co-aggregation of p62, and there may be structural differences between mutant and wildtype.

In the GCL, two of the four *UBQLN2*-linked ALS/FTD cases were devoid of pTDP-43, ubiquilin 2, and p62 aggregates, consistent with previous findings [1,38] (case P497H, Fig. 4A_1_; and MN17 – T487I, Fig. 4B_1_). In contrast, case V:7 – T487I showed very sparse pTDP-43 cytoplasmic aggregates in the GCL that were negative for ubiquilin 2 but weakly positive for p62 (Fig. 4C_1_, white hollow arrowhead). Consistent with Gkazi and colleagues [15] and with the known spectrum of pTDP-43 loads in the GCL in ALS/FTD [55], the *UBQLN2* p.P506S case had abundant p62-positive pTDP-43 cytoplasmic aggregates in the GCL, that either co-localised with ubiquilin 2 (Fig. 4D_1_, white filled arrowheads) or were independent of ubiquilin 2 (Fig. 4D_1_, white hollow arrowheads). Mutant ubiquilin 2 aggregation in the GCL was therefore likely scaffolded or ‘seeded’ by pTDP-43 when present. The *UBQLN2* p.S222G PSP case showed sparse p62-only aggregates in the GCL that were not seen in *UBQLN2*-linked ALS/FTD cases (Fig. 4E_1_, pink arrowheads).

*UBQLN2-*linked ALS/FTD case P506S also had sparse pTDP-43 aggregates in the CA – pyr cells, which were positive for both ubiquilin 2 and p62 (Fig. 4D_3_, white filled arrowhead). All *UBQLN2* variant and mutation cases were negative for DPRs (not shown).

#### Combined neuropathological signatures discriminate between *UBQLN2*-linked, *C9orf72*-linked, sporadic, and familial cases

Integration of all neuropathological findings (Fig. 5) revealed a characteristic hippocampal neuropathological signature for *UBQLN2*-linked ALS/FTD, that was distinct from that in other forms of ALS/FTD. Sporadic ALS cases were wholly devoid of hippocampal ubiquilin 2 or DPR protein pathology, with a minority of cases showing pTDP-43 aggregates in the granule cells that were ubiquilin 2 negative. Thus, wildtype ubiquilin 2 is not seeded/ scaffolded to aggregate by pTDP-43 aggregation. *C9orf72*-linked cases showed ML ubiquilin 2 aggregates that were wispy or punctate and predominantly p62 negative, while *UBQLN2*-linked cases showed ML ubiquilin 2 aggregates that were punctate and predominantly p62 positive. Mutant ubiquilin 2 aggregates thus promote the co-aggregation of p62, and this may relate to their conformational differences. Overall, mutant ubiquilin 2 causes unique neuropathology that is shared by p.P506S, p.P497H, and p.T487I ubiquilin 2 (Fig. 6). The absence of this signature in a case harbouring a p.S222G variant manifesting with PSP suggests this neuropathology can be used to discriminate pathological ALS/FTD-causing *UBQLN2* mutations.

**Figure 5.**
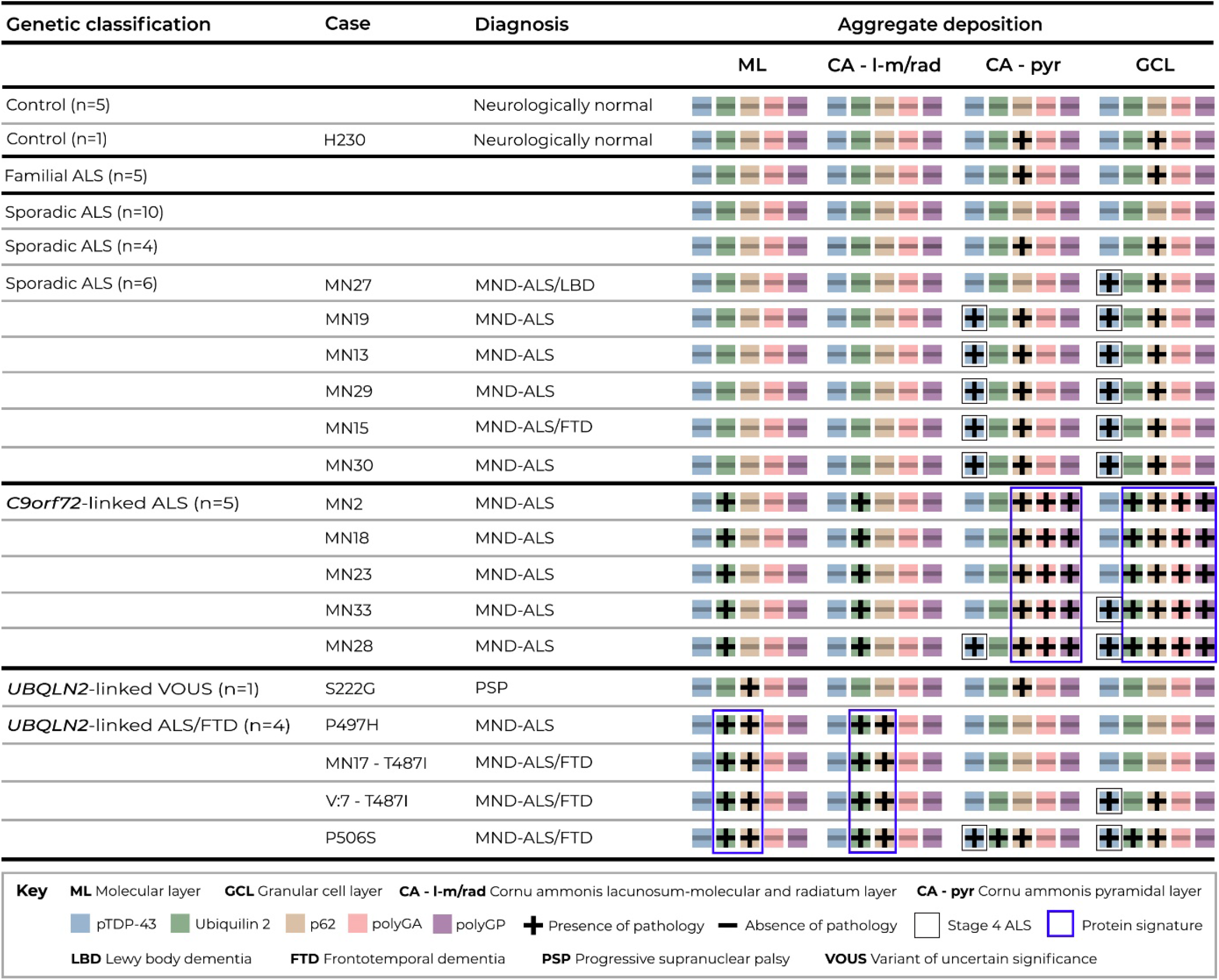
Combined pTDP-43, ubiquilin 2, p62, and dipeptide repeat protein immunohistochemical staining discriminated between sporadic, *C9orf72*-linked, and *UBQLN2*-linked ALS/FTD. Key shown within figure. P62 aggregates (‘+’ symbol on brown box) without other co-labelling were observed in CA – pyr and GCL cells in a subset of neurologically normal controls (n=1 of 5), unrelated familial ALS (n=5 of 5), and sporadic ALS cases (n=4 of 20). In contrast, pTDP-43 pathology (boxed ‘+’ symbol on blue box) was present in these layers in a separate subset of sALS cases (n=6 of 20), co-localising with p62. Ubiquilin 2 pathology (‘+’ symbol on green box) was present in the hippocampus when ubiquilin 2 was wildtype (*C9orf72*-linked ALS) or mutant (*UBQLN2*-linked ALS/FTD). Wildtype ubiquilin 2 in *C9orf72*-linked ALS ML and CA – lm/rad regions was p62-negative, and in the CA – pyr layer was associated with polyGA, polyGP, and p62 aggregates. PolyGA and polyGP aggregates were found in the GCL with or without ubiquilin 2, all colocalising with p62. CA – pyr and GCL aggregates had additional pTDP-43 pathology in only one case, and even then, only rarely. Mutant ubiquilin 2 aggregates in *UBQLN2*-linked ALS/FTD ML, and CA – lm/rad regions were p62-positive and associated with GCL pTDP-43 aggregates only in some cases and either with or without ubiquilin 2 co-labelling. Blue outlines indicate unique aggregation features of *C9orf72*-linked ALS and *UBQLN2*-linked ALS/FTD hippocampal pathology. Abbreviations: CA – lm/rad, cornu ammonis – lacunosum-molecular and radiatum layers; CA – pyr, cornu ammonis – pyramidal cells; GCL, granule cell layer.

**Figure 6.**
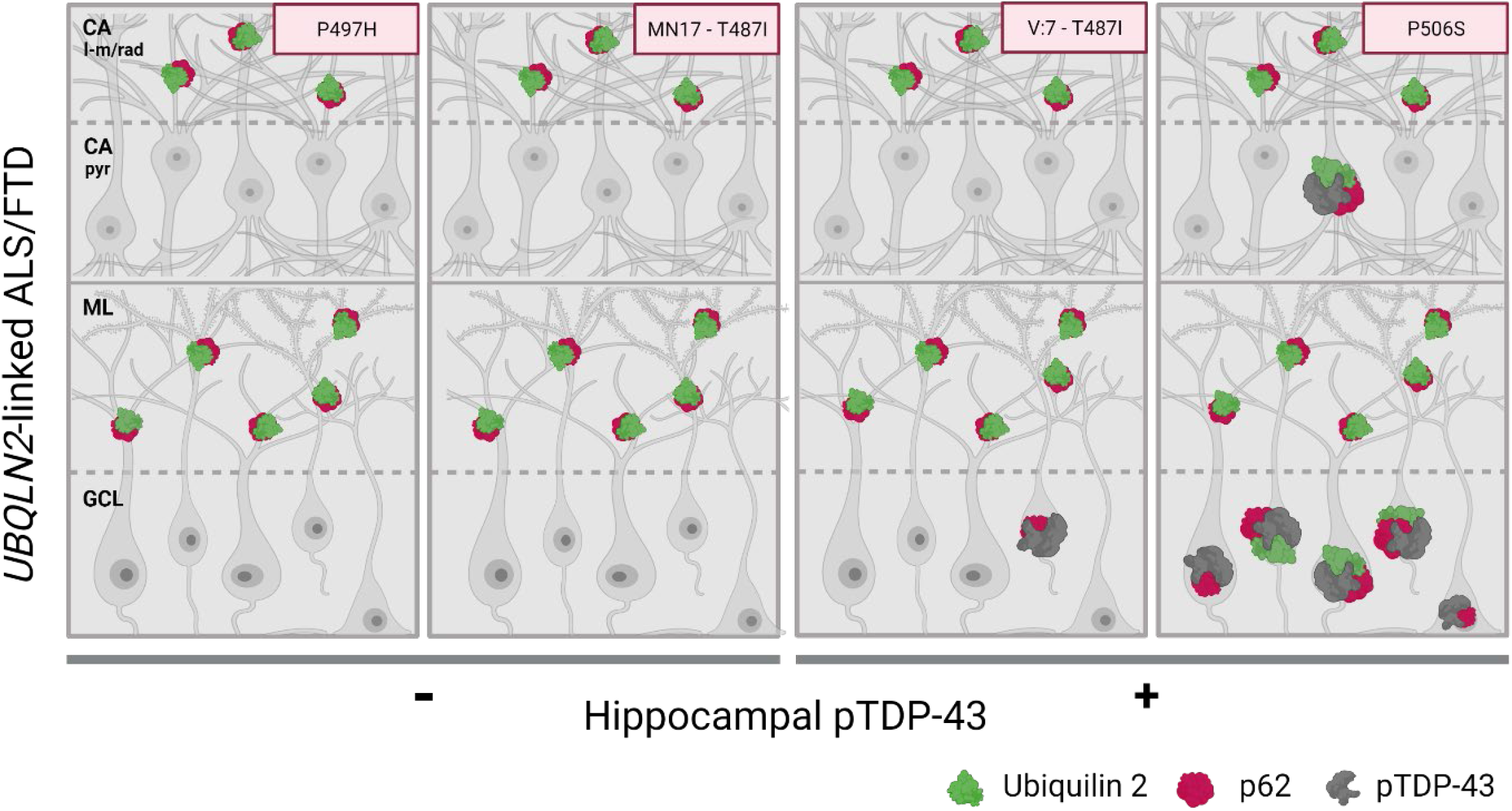
Schematic representation of the hippocampal neuropathological signature defining *UBQLN2*-linked ALS/FTD. In *UBQLN2*-linked ALS/FTD cases without hippocampal pTDP-43 proteinopathy, mutant ubiquilin 2 is punctate, p62-positive, and forms aggregates exclusively in the CA l-m/rad layer and ML (cases p.P497H and MN17 – p.T487I). In *UBQLN2*-linked ALS/FTD cases with hippocampal pTDP-43 proteinopathy, in addition to punctate p62-positive ubiquilin 2 aggregates in the CA l-m/rad, and molecular layers, there are granule cell layer pTDP-43 aggregates that are either; frequent and ubiquilin 2-labelled (case p.P506S), or rare and ubiquilin 2-negative (case V7-p.T487I), suggesting a pathological cascade in which granule cell layer pTDP-43 aggregates provide a scaffold around which mutant ubiquilin 2 can aggregate. Abbreviations: CA – lm/rad, cornu ammonis – lacunosum-molecular and radiatum layers; CA – pyr, cornu ammonis – pyramidal cells; GCL, granule cell layer. Image created in Biorender.com

Ubiquilin 2 hippocampal pathology was present in both *UBQLN2*-linked ALS/FTD and *C9orf72*-linked ALS, but these genotypes could be discriminated when ubiquilin 2 was co-labelled with pTDP-43, DPR proteins, or p62. *C9orf72*-linked cases always showed GCL ubiquilin 2 that co-localised with DPR aggregates, but rarely pTDP-43. This supports the lack of seeding of wildtype ubiquilin 2 aggregation by pTDP-43 but suggests that wildtype ubiquilin 2 can be seeded by polyGA. In contrast, *UBQLN2*-linked cases showed GCL ubiquilin 2 if co-localised pTDP-43 was present. Therefore, mutant ubiquilin 2 is more aggregation-prone than wildtype, being seeded by pTDP-43.

#### *UBQLN2* p.T487I mutation in ALS/FTD families FALS5 and FALS14 was inherited from a common ancestor

Since the initial report by Williams et al. [17] of an identical *UBQLN2* p.T487I mutation in ALS families FALS5 and FALS14, we report here that cases MN17 (IV:18) and V:7 from family FALS5 developed ALS+FTD, indicating that FTD is part of the clinical phenotype in that family. To further examine relatedness between the families and confirm that the *UBQLN2* p.T487I mutation arose in a common founder, we performed identity-by-descent (IBD) analysis. IBD segments were identified over the *UBQLN2* locus between all four affected individuals from both families (Table 1), while there were no IBD segments inferred over *UBQLN2* between the affected individuals from FALS5 and the unaffected individual from FALS14 who did not carry the *UBQLN2* p.T487I mutation. The interval shared by all four affected individuals spanned rs952836 to rs6423133 and is 68 cM in length. This confirms a founder effect of *UBQLN2* p.T487I in FALS5 and FALS14. The genotyped individuals across these families are estimated to be 4th-to 5th-degree relatives (1st cousins once removed -2nd cousins) (Table 2), now confirming segregation of the *UBQLN2* p.T487I mutation in 17 individuals from the proposed combined Australia-New Zealand pedigree and providing further strong genetic evidence, in addition to the neuropathological evidence, that *UBQLN2* p.T487I is pathogenic for ALS/FTD.

**Table 1.** IBD segments inferred over *UBQLN2* on the X chromosome (chromosome 23).

| Individual 1 |  | Individual 2 |  |  |  |  |  |
| --- | --- | --- | --- | --- | --- | --- | --- |
| Family ID | Pedigree ID | Family ID | Individual ID | StartSNP | EndSNP | Start Pos(bp) | End Pos(bp) |
| FALS5 | III:8 | FALS5 | IV:9 | rs17330993 | rs11156600 | 2779749 | 154440161 |
| FALS5 | III:8 | FALS5 | IV:18<sup>a</sup> | rs7062445 | rs6423133 | 19650411 | 123608292 |
| FALS5 | III:8 | FALS14 | III:2 | rs952836 | rs17315029 | 40344087 | 129605268 |
| FALS5 | IV:9 | FALS5 | IV:18 | rs6628597 | rs6423133 | 31382037 | 123608292 |
| FALS5 | IV:9 | FALS14 | III:2 | rs952836 | rs17277770 | 40344087 | 132774456 |
| FALS5 | IV:18 | FALS14 | III:2 | rs952836 | rs6423133 | 40344087 | 123608292 |

<sup>a</sup> FALS5 IV:18 is referred to as MN17 in this report.

**Table 2.** Estimated degree of relatedness and fraction of genome with zero, one, and two alleles identical-by-descent (IBD) between all samples in *UBQLN2* p.T487I-linked families FALS5 and FALS14.

|  | Individual 1 |  | Individual 2 |  | Fraction IBD=0 | Fraction IBD=1 | Fraction IBD=2 | Degree<sup>a</sup> |
| --- | --- | --- | --- | --- | --- | --- | --- | --- |
|  | Family ID | Pedigree ID | Family ID | Individual ID |  |  |  |  |
| Intra-family | FALS5 | III:8 | FALS5 | IV:9 | 0.529 | 0.471 | 0 | 2 |
|  | FALS5 | III:8 | FALS5 | IV:18<sup>b</sup> | 0.529 | 0.471 | 0 | 2 |
|  | FALS5 | IV:9 | FALS5 | IV:18 | 0.751 | 0.249 | 0 | 3 |
|  | FALS14 | II:1 | FALS14 | III:2 | 0.515 | 0.485 | 0 | 2 |
| Inter-family | FALS5 | III:8 | FALS14 | II:1 | 0.877 | 0.123 | 0 | 4 |
|  | FALS5 | III:8 | FALS14 | III:2 | 0.927 | 0.073 | 0 | 5 |
|  | FALS5 | IV:9 | FALS14 | II:1 | 0.942 | 0.058 | 0 | 5 |
|  | FALS5 | IV:9 | FALS14 | III:2 | 0.948 | 0.052 | 0 | 5 |
|  | FALS5 | IV:18 | FALS14 | II:1 | 0.926 | 0.074 | 0 | 5 |
|  | FALS5 | IV:18 | FALS14 | III:2 | 0.948 | 0.052 | 0 | 5 |

<sup>a</sup> Degrees of relatedness as per [100]:
2 – Grandparent-grandchild, Avuncular, Half-sibling
3 – First cousin, Great-grandparent, Grand-avuncular
4 – First cousin once removed, GG-grandparent
5 – Second cousin, First cousin twice removed
<sup>b</sup> FALS5 IV:18 is referred to as MN17 in this report.

## Discussion

ALS shows considerable clinical, pathological, and genetic heterogeneity [58–60]. While TDP-43 proteinopathy is seen in 97% of cases [40,43,61], ALS/FTD-causing genetic mutations can cause deposition of the encoded mutant protein leading to additional pathological aggregate signatures. We previously exploited this to infer *C9orf72* repeat expansion genotypes [62,63] through neuropathological screening of our New Zealand ALS cases [38]. Mutant *SOD1* [64,65], *FUS* [44], and certain other genotypes [66,67] can also be inferred from neuropathology. However, for many ALS/FTD genes — particularly those such as *TARDBP*, *SQSTM1*, and *UBQLN2* that encode proteins already within the hallmark TDP-43 inclusions — no completely discriminating neuropathology has been reported [42,66,68–72]. This has hampered the validation of pathogenicity of novel variants in these genes, in turn obscuring understanding of protein domains and molecular processes important to ALS/FTD pathogenesis. However, we report here a unique neuropathological signature for mutant ubiquilin 2 in the hippocampus that discriminates *UBQLN2*-linked ALS/FTD cases from all others tested.

### pTDP-43 in *UBQLN2*-linked cases: An independent pathology

Before discussing the unique pattern of hippocampal ubiquilin 2 pathology shared by *UBQLN2*-linked cases, it must first be noted that pTDP-43 deposition was variable. Two of the four *UBQLN2-*linked cases were devoid of GCL pTDP-43 aggregates (p.P497H (ALS) and MN17-p.T487I (ALS+FTD)), one case had very sparse GCL pTDP-43 aggregates that were ubiquilin 2-negative (V:7-p.T487I (ALS+FTD)), and one case showed frequent GCL pTDP-43 aggregates that were mostly ubiquilin 2-positive (p.P506S (ALS+FTD)). Although pTDP-43 aggregates are nearly ubiquitous in the ALS spinal cord and motor cortex, only in ∼15-30% of ALS cases are they found in the hippocampus [55,73]. Our findings suggest that variable regional pTDP-43 deposition occurs in the context of *UBQLN2*-linked ALS/FTD just as it does in sporadic and *C9orf72*-linked ALS/FTD.

### Wildtype ubiquilin 2 pathology in *C9orf72*-linked cases

Hippocampal ubiquilin 2 deposition is a known and striking feature of *C9orf72*-linked ALS/FTD [39]. In our *C9orf72*-linked ALS cases, large stellate GCL ubiquilin 2 was found to preferentially co-localise with the aggregation-prone polyGA DPR protein compared to polyGP. Furthermore, polyGP aggregates were very rarely independent of polyGA. These findings support the emerging consensus that polyGA aggregates ‘seed’ polyGP protein aggregation [74–76]. Similar observations were made by Mackenzie and colleagues [56] of aggregates with a core of polyGA surrounded by an aggregated TDP-43 shell; a finding recapitulated here and supported by *in vitro* work showing polyGA aggregation preceding TDP-43 accumulation [77]. The ability of DPR proteins to seed ubiquilin 2 however, appears more complex. We previously showed ubiquilin 2 at the core of polyGP-positive aggregates, but we did not co-label for polyGA [38]. STED imaging in the current study shows that ubiquilin 2 may either surround polyGA or be enmeshed with it, suggesting that in *C9orf72-*linked ALS pathogenesis, the interaction between aggregating polyGA and ubiquilin 2 may be an early event.

In contrast to ubiquilin 2 co-aggregation with large stellate DPR proteins in the GCL, small neuritic ubiquilin 2 aggregates in *C9orf72*-linked cases punctuated the ML, and CA – lm/rad regions seemingly independent of a nucleating protein. The neurites in which these small aggregates are found likely derive from the granule or pyramidal cells, respectively [78,79], so DPR protein inclusions in the somata may promote the aggregation of ubiquilin 2 in the dendrites of the same cell. DPR aggregates can sequester proteasome components [80–82], and loss of C9orf72 protein function in cells expressing the DPR-encoding variant 2 [83] can impair autophagy [84–89], which may underpin wildtype ubiquilin 2 aggregation in the ML, and CA – lm/rad layers in *C9orf72*-linked cases.

### Mutant ubiquilin 2 pathology in *UBQLN2*-linked cases: A neuropathological signature

Even in its wildtype state, ubiquilin 2 intrinsically self-assembles [3] but there is now ample biophysical evidence demonstrating that mutations to ubiquilin 2 confer an increased propensity to oligomerise and undergo aberrant LLPS, forming insoluble aggregates within the cell [8,9,12,90–92]. Here we confirm that mutant ubiquilin 2 in the human hippocampal ML (granule cell dendrites) is aggregated under less permissive conditions than wildtype ubiquilin 2, appearing to require no aggregated protein scaffold, or protein aggregation event in the GCL (granule cell soma).

Our study additionally found that p62 labelling was required, while pTDP-43 labelling was dispensable, to confirm cases as being *UBQLN2*-linked. P62 co-localises with pTDP-43 aggregates and DPR proteins in ALS/FTD [62,93,94], or with hyperphosphorylated tau in a range of tauopathies [95–97], or with mutant α-synuclein in synucleinopathies [97,98]. Given this promiscuity for substrates, discrimination between mutant and wildtype ubiquilin 2 by p62 suggests that there are unique structural or biochemical features of mutant ubiquilin 2 aggregates. This leads to the intriguing possibility that mutant ubiquilin 2 would be selectively druggable.

In addition to mechanistic insights, the mutant ubiquilin 2 neuropathological signature we describe will enable classification of *UBQLN2* variants of uncertain significance, clarifying the implications of a positive genetic result in a patient. Currently, only four missense mutations in *UBQLN2* are designated by ClinVar as pathogenic or likely pathogenic (resulting in p.M392V, p.Q425R, p.P497H, p.P497S, p.P506T, https://www.ncbi.nlm.nih.gov/clinvar/, accessed July 11 2023 [99]). However, 62 other *UBQLN2* missense changes listed in ClinVar are classified as benign or of uncertain significance, including c.1516C>T resulting in p.P506S. Indeed, neither *UBQLN2* p.T487I nor p.S222G are even listed in ClinVar, leaving individual diagnostics labs to perform classification for patient reporting. We now confirm that two ALS/FTD cases with p.T487I, which segregates with disease in a large kindred and shows early onset in males consistent with X-linked disease, shared our newly identified mutant ubiquilin 2 neuropathological signature, supporting its classification as pathogenic. Conversely, a male PSP case with p.S222G who lived to 93 y did not share this signature, suggesting this variant is benign. We encourage uptake of this hippocampal ubiquilin 2 neuropathology signature, by other labs or in collaboration with the authors hereof, as a tool to explore *UBQLN2* variant pathogenicity.

## Conclusion

Ubiquilin 2 aggregates were seen in the hippocampus of ALS/FTD cases across a range of genotypes. Wildtype ubiquilin 2 *in vitro* is known to be aggregation-prone; in human brain it co-aggregated with polyGA, but not pTDP-43 and with little co-aggregation of p62. Mutant ubiquilin 2 *in vitro* is known to be more aggregation-prone than wildtype; in brain it was either co-aggregated with pTDP-43 or aggregated independently of a known scaffold and appeared to promote the co-aggregation of p62. This hippocampal ubiquilin 2 neuropathology signature demonstrates that ubiquilin 2 aggregation is likely to play a mechanistic role in *C9orf72*-linked and *UBQLN2*-linked ALS/FTD and provides a definitive framework for exploring the biological implications of *UBQLN2* genetic variation.

### List of abbreviations

ALS: Amyotrophic lateral sclerosis
bvFTD: Behavioural variant FTD
CA-lm/rad: Cornu ammonis lacunosum-molecular and radiatum layers
CA-pyr: Cornu ammonis pyramidal layer
C9orf72: Chromosome 9 open reading frame 72
DPR: Dipeptide repeat
FFPE: Formalin-fixed paraffin-embedded
FTD: Frontotemporal dementia
GCL: Granule cell layer
IBD: Identity by descent
LLPS: Liquid-liquid phase separation
ML: Molecular layer
NCI: Neuronal cytoplasmic inclusions
PBS: Phosphate-buffered saline
PolyGA: Poly(glycine-arginine)
PolyGP: Poly(glycine-proline)
pTDP-43: Phosphorylated TDP-43
SOD1: Superoxide dismutase
STED: Stimulated emission depletion
SQSTM1: Sequestosome 1
TDP-43: Transactive response DNA binding protein 43 kDa

## Disclosures and declarations

### Data transparency

The datasets used and/or analysed during the current study are available from the corresponding author on reasonable request.

## Compliance with ethical standards

### Research involving human participants

All protocols were approved by the University of Auckland Human Participants Ethics Committee (New Zealand) and carried out as per approved guidelines. This study was also approved by the Human Research Ethics Committee of Macquarie University (520211013428875).

### Informed consent

Informed donor consent and ethical approvals were obtained at each site as described previously [1,15,17,38].

## Acknowledgements and funding

This publication is dedicated to the incredible patients and families who contribute to our research. We thank Marika Eszes at the Centre for Brain Research, University of Auckland, New Zealand; Nailah Siddique at the Northwestern University Feinberg School of Medicine, Chicago, USA; Sashika Selvaduncko and Claire Troakes at the London Neurodegenerative Diseases Brain Bank and Brains for Dementia; and the Neurological Foundation of New Zealand for their ongoing financial support of the Human Brain Bank. We also thank Fairlie Hinton and Dr. Catriona McLean at the Victorian Brain Bank, which is supported by The Florey Institute of Neuroscience and Mental Health, The Alfred and the Victorian Forensic Institute of Medicine and funded in part by Parkinson’s Victoria, MND Victoria, FightMND and Yulgilbar Foundation. The imaging data reported in this paper were obtained at the Biomedical Imaging Research Unit (BIRU), operated by the Faculty of Medical and Health Sciences’ Technical Services at the University of Auckland. KT was funded by a doctoral scholarship from Amelia Pais-Rodriguez and Marcus Gerbich. BVD and MD were funded by the Michael J Fox Foundation [Grant ID: 16420]. BVD was also funded by a Health Research Council Sir Charles Hercus Health Research Fellowship [21/034]. ELS was supported by Marsden FastStart and Rutherford Discovery Fellowship funding from the Royal Society of New Zealand [15-UOA-157, 15-UOA-003]. This work was also supported by grants from Motor Neuron Disease NZ, Freemasons Foundation of New Zealand, Matteo de Nora, Sir Thomas and Lady Duncan Trust and the Coker Family Trust (to MD), and PaR NZ Golfing. No funding body played any role in the design of the study, nor in the collection, analysis, or interpretation of data nor in writing the manuscript.

