## Supplementary Material for "Hippocampal protein aggregation signatures fully distinguish pathogenic and wildtype *UBQLN2* in amyotrophic lateral sclerosis"

### Supplementary figures and tables

Kyrah M. Thumbadoo[1,2], Birger V. Dieriks[2,3], Helen C. Murray[2,3], Molly E. V. Swanson[1,3], Ji Hun Yoo[2,4], Nasim F. Mehrabi[2,3], Clinton Turner[2,3,5], Michael Dragunow[2,4], Richard L.M. Faull[2,3], Maurice A. Curtis[2,3], Teepu Siddique[6], Christopher E Shaw[2,7], Lyndal Henden[8], Kelly L. Williams[8], Garth A. Nicholson[8,9,10,11], Emma L. Scotter[1,2]

1 School of Biological Sciences, University of Auckland, Auckland, New Zealand

2 Centre for Brain Research, University of Auckland, Auckland, New Zealand

3 Department of Anatomy and Medical Imaging, University of Auckland, Auckland, New Zealand

4 Department of Pharmacology and Clinical Pharmacology, University of Auckland, Auckland, New Zealand

5 Department of Anatomical Pathology, LabPlus, Auckland City Hospital, Auckland, New Zealand

6 Departments of Neurology, Cell and Developmental Biology and Pathology, Northwestern University Feinberg School of Medicine, Chicago, USA

7 UK Dementia Research Institute Centre, Institute of Psychiatry, Psychology and Neuroscience, King's College London, United Kingdom

8 Macquarie University Centre for Motor Neuron Disease Research, Macquarie Medical School, Faculty of Medicine, Health and Human Sciences, Macquarie University, Sydney, New South Wales, Australia

9 Northcott Neuroscience Laboratory, Australian and New Zealand Army Corps (ANZAC) Research Institute, Concord, New South Wales, Australia

10 Faculty of Medicine, University of Sydney, Sydney, New South Wales, Australia

11 Molecular Medicine Laboratory, Concord Repatriation General Hospital, Concord, New South Wales, Australia

**Supplementary Table 1. Demographics and clinical features of study cohort**

| Case type | UoA case code | Diagnosis | Sex | AAD (y) | PMD (h) | Genetic mutation |
|---|---|---|---|---|---|---|
| **Control** | H211 | NNDC | M | 41 | 9.5 | N/A |
| | H215 | NNDC | F | 67 | 23.5 | N/A |
| | H230 | NNDC | F | 57 | 32 | N/A |
| | H238 | NNDC | F | 63 | 16 | N/A |
| | H239 | NNDC | M | 64 | 15.5 | N/A |
| | H247 | NNDC | M | 51 | 31 | N/A |
| **Sporadic ALS** | MN4 | MND-ALS | M | 41 | 7 | None found[a] |
| | MN5 | MND-ALS | F | 55 | 5 | None found[a] |
| | MN6 | MND-ALS | M | 58 | 8.5 | None found[a] |
| | MN8 | MND-ALS | M | 84 | 3 | None found[a] |
| | MN9 | MND-ALS | M | 88 | 36 | None found[a] |
| | MN10 | MND-ALS | M | 46 | >24 | None found[a] |
| | MN12 | MND-ALS | M | 49 | 34 | None found[a] |
| | MN13 | MND-ALS | M | 55 | 10 | None found[a] |
| | MN15 | MND-ALS/FTD | F | 54 | 18 | None found[a] |
| | MN16 | MND-ALS | M | 69 | 16.5 | None found[a] |
| | MN19 | MND-ALS | M | 75 | 20.5 | None found[a] |
| | MN20 | MND-ALS | M | 85 | 15 | None found[a] |
| | MN22 | MND-ALS | F | 65 | 9 | ND |
| | MN25 | MND-ALS | M | 71 | 23.5 | ND |
| | MN26 | MND-ALS | M | 77 | 3 | ND |
| | MN27 | MND-ALS+LBD-bs | F | 87 | - | ND |
| | MN29 | MND-ALS | M | 72 | 24 | ND |
| | MN30 | MND-ALS | F | 84 | 17.5 | ND |
| | MN31 | MND-ALS | F | 58 | 19 | ND |
| | MN32 | MND-ALS | M | 59 | 6 | ND |
| **Familial ALS of unknown genotype** | MN11 | MND-ALS | F | 77 | 18 | None found[a] |
| | MN14 | Prob-ALS | F | 59 | 19 | None found[a] |
| | MN21 | MND-ALS | F | 59 | 17.5 | None found[a] |
| ***SOD1*-linked** | MN24 | MND-ALS | F | 54 | - | *SOD1* p.E101G |
| ***FUS*-linked** | A203/12 | MND-ALS | F | 23 | 37 | *FUS* p.P525L |
| ***UBQLN2* VOUS or *UBQLN2*-linked** | S222G | PSP | M | 93 | 19.5 | *UBQLN2* p.S222G |
| | MN17 – T487I (FALS5 IV:18) | MND-ALS/FTD | F | 61 | 69 | *UBQLN2* p.T487I |
| | V:7 – T487I (FALS5 V:7) | MND-ALS/FTD | M | 39 | 25.5 | *UBQLN2* p.T487I |
| | P497H | ALS | F | 57 | - | *UBQLN2* p.P497H |
| | P506S | MND-ALS/FTD | F | 56 | 35 | *UBQLN2* p.P506S |
| ***C9orf72*-linked** | MN2 | MND-ALS | F | 53 | >24 | *C9orf72* repeat expansion[b] |
| | MN18 | MND-ALS | F | 53 | 12 | *C9orf72* repeat expansion[c] |
| | MN23 | MND-ALS | F | 79 | 27 | *C9orf72* repeat expansion[d] |

| Case type | UoA case code | Diagnosis | Sex | AAD (y) | PMD (h) | Genetic mutation |
|---|---|---|---|---|---|---|
| | MN28 | MND-ALS | F | 62 | 14 | *C9orf72* repeat expansion[e] |
| | MN33 | MND-ALS | M | 65 | 46 | *C9orf72* repeat expansion[d] |

[a] No mutations detected in *C9orf72, TARDBP, FUS,* or *SOD1,* as previously described in [2].

[b] Obligate carrier, affected offspring genotyped.

[c] Genotyped in [2].

[d] Genotyped during life.

[e] Genotype inferred from neuropathology.

Abbreviations: -, unknown; AAD, age at death; LBD-bs, Lewy body disease- brainstem predominant; N/A, not applicable; ND, not done; NNDC, non-neurologically diseased control; PMD, post-mortem delay; PSP, progressive supranuclear palsy .

### Supplementary Table 2. Primary antibodies

| Primary antibody | Species (isotype) | Manufacturer | Catalogue # | RRID | Clonality | Dilution |
|---|---|---|---|---|---|---|
| **Multiplex Fluorescent Immunohistochemistry panel** | | | | | | |
| C9RANT (PolyGA) | Mouse (IgG1κ) | Merck Millipore | MABN889 | AB_2728663 | Monoclonal | 1:2000 |
| Ubiquilin 2 | Mouse ($IgG_2a$) | Santa Cruz Biotechnology | SC-100612 | AB_2272422 | Monoclonal | 1:1000 |
| C9RANT (PolyGP) | Rabbit (IgG) | Novus Biologicals | NBP2-25018 | AB_2893239 | Polyclonal | 1:3000 |
| pTDP-43 (Ser409/410) | Rat ($IgG_2a$) | BioLegend | BL829901, Clone 1D3 | AB_2564934 | Monoclonal | 1:3000 |
| p62 | Guinea Pig (IgG) | Progen | GP62-C | AB_2687531 | Polyclonal | 1:500 |
| **Double-label Fluorescent Immunohistochemistry for STED imaging panel** | | | | | | |
| C9RANT (PolyGA) | Mouse (IgG1κ) | Merck Millipore | MABN889 | AB_2728663 | Monoclonal | 1:2000 |
| Ubiquilin 2 | Mouse (IgG2a) | Santa Cruz Biotechnology | SC-100612 | AB_2272422 | Monoclonal | 1:1000 |

Abbreviations: RRID, Research resource identifier; pTDP-43, phosphorylated (Ser409/410) TDP-43

### Supplementary Table 3. Secondary antibodies and stains

| Secondary antibody | Conjugate | Manufacturer | Catalogue # | RRID | Clonality | Dilution |
|---|---|---|---|---|---|---|
| **Multiplex Fluorescent Immunohistochemistry panel** | | | | | | |
| Goat anti-mouse IgG1κ | Alexa Fluor® 488 | Thermo Fisher Scientific | A-21121 | AB_2535764 | Monoclonal | 1:500 |
| Goat anti-mouse $IgG_2a$ | Alexa Fluor® 594 | Thermo Fisher Scientific | A-21135 | AB_2535774 | Monoclonal | 1:500 |
| Goat anti-rabbit IgG | IRDye 800CW | LI-COR Biosciences | 926-32211 | AB_2651127 | Polyclonal | 1:500 |
| Goat anti-rat $IgG_2a$ | Alexa Fluor® 546 | Thermo Fisher Scientific | A-11081 | AB_141738 | Polyclonal | 1:500 |
| Goat anti-guinea pig IgG | Alexa Fluor® 647 | Thermo Fisher Scientific | A-21450 | AB_141882 | Polyclonal | 1:500 |
| **Stain** | **Species (isotype)** | **Company** | **Catalogue #** | **RRID** | **Clonality** | **Dilution** |
| Hoechst 33342 nuclear stain | - | Thermo Fisher Scientific | H3750 | | - | 1:2000 |
| **Double-label Fluorescent Immunohistochemistry for STED imaging panel** | | | | | | |
| Goat anti-mouse IgG1k | Biotin | Thermo Fisher Scientific | A10519 | AB_1500809 | Monoclonal | 1:500 |
| Abberior Star Red | Neutravidin | Abberior | STRED-0121 | | - | 1:500 |
| Goat anti-mouse $IgG_2a$ | Alexa Fluor® 594 | Thermo Fisher Scientific | A-21135 | AB_2535774 | Monoclonal | 1:500 |

Abbreviations: RRID, Research resource identifier; pTDP-43, phosphorylated (Ser409/410) TDP-43

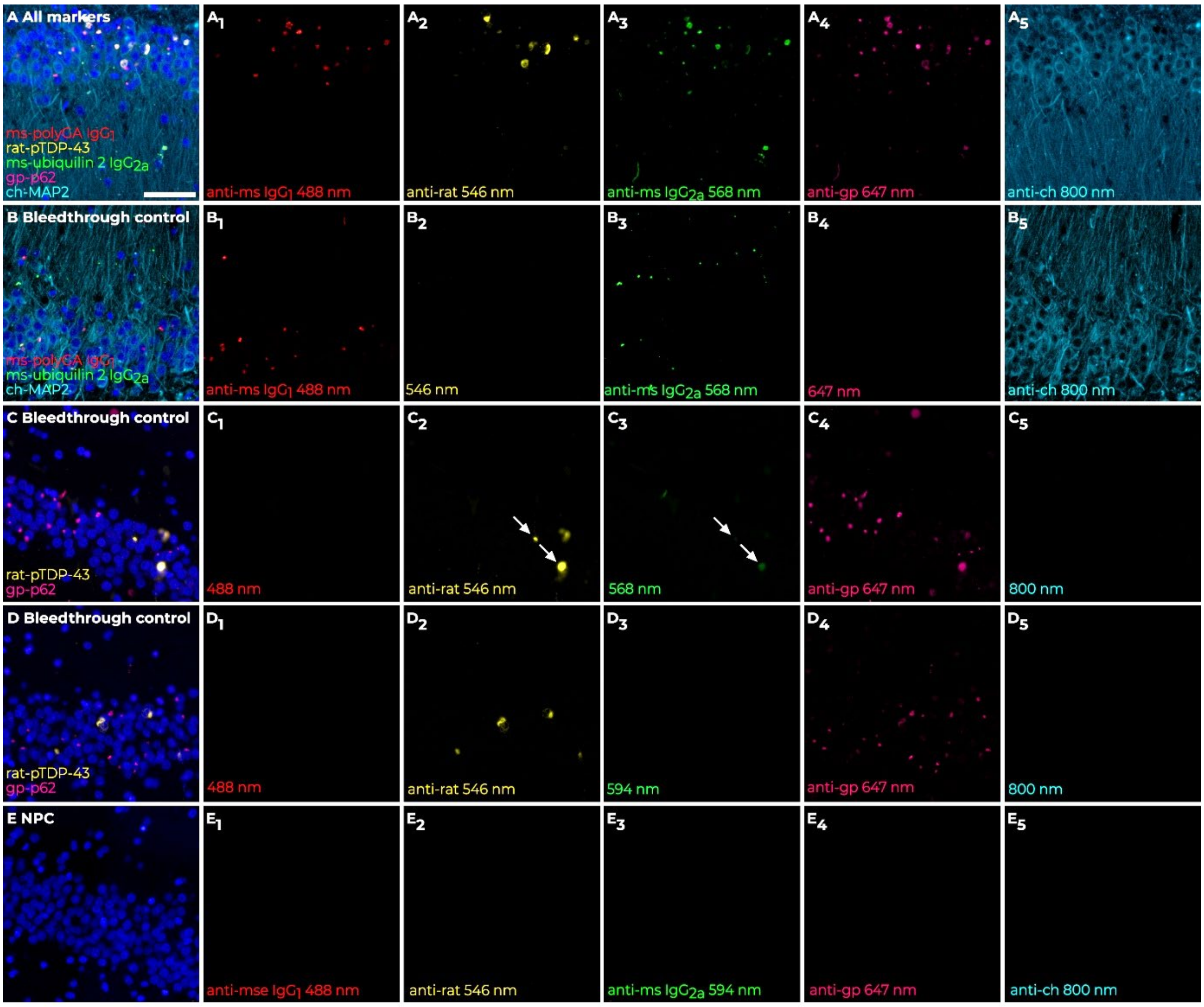


**Supplementary Figure 1. Multiplex immunohistochemistry secondary antibody bleedthrough, bleedback, and secondary-only controls.** Aggregate staining was first optimised in *C9orf72*-linked ALS case MN28, as it stains for all markers (**A-A5**). DPR protein polyGA staining in the 488 nm channel (**B1**) demonstrated no bleedthrough into the next channel, 546 nm (**B2**). Additionally, ubiquilin 2 staining in 568 nm (B3) showed no bleedback into 546 nm (**B2**), nor bleedthrough into the 647 nm channel (**B4**). pTDP-43 staining in the 546 nm channel (**C2**) showed trace amounts of bleedthrough into the 568 nm channel (**C3**, white arrowheads). As a result, a 594 nm secondary antibody was used to detect ubiquilin 2 instead and when imaged, showed no bleedthrough of pTDP-43 (**D3**). Secondary antibodies were specific to primary antibodies directed against polyGA, pTDP-43, ubiquilin 2, p62, and MAP2 as no staining was observed when primary antibodies were omitted (**E-E5**). Scale bar for all images, 50 µm

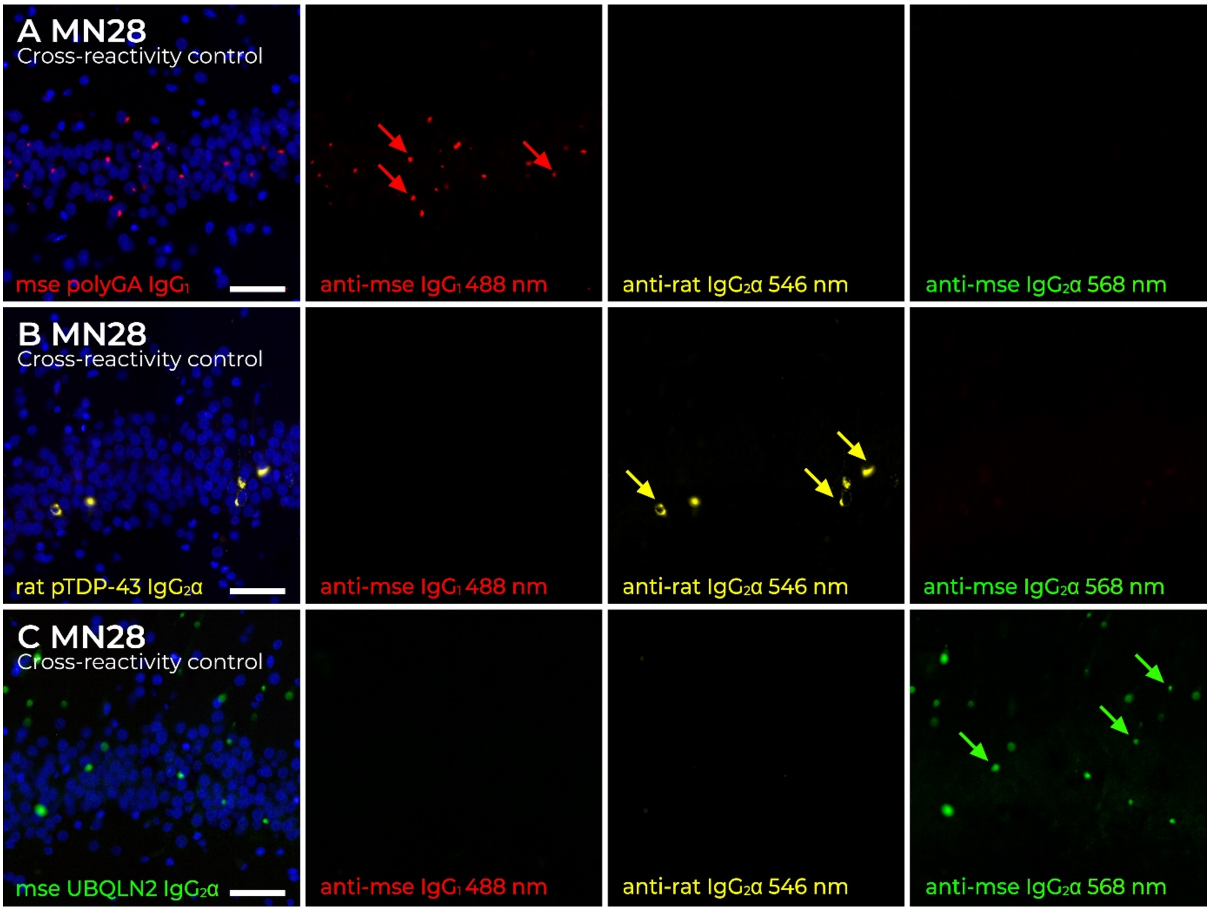


**Supplementary Figure 2. Multiplex immunohistochemistry secondary antibody cross-reactivity controls.** Aggregate staining was first optimised in *C9orf72*-linked ALS case MN28, as it stains for all markers. DPR protein polyGA staining using a mouse IgG1 488 secondary (**A**, red arrows) was not detected by secondaries rat IgG2a 546 nm and mouse IgG2a 568 nm when all secondaries were added. Additionally, pTDP-43 staining in 546 nm (**B**, yellow arrows) was not detected by mouse secondaries IgG1 488 nm and IgG2a 568 nm. Finally, a mouse IgG2a secondary specifically stained for ubiquilin 2 aggregates (**C**, green arrows), as no staining was observed in the 488 nm and 546 nm channels. Note: In the final immunohistochemical runs for all cohort cases, a mouse IgG2a 594 secondary was used to visualise ubiquilin 2 staining. The 568 secondary antibody shown in this optimisation and the 594 secondary antibody are equivalent as per manufacturers information, with different conjugated fluorophores. Thus, these cross-reactivity tests are still valid. Scale bar for all images, 50 µm

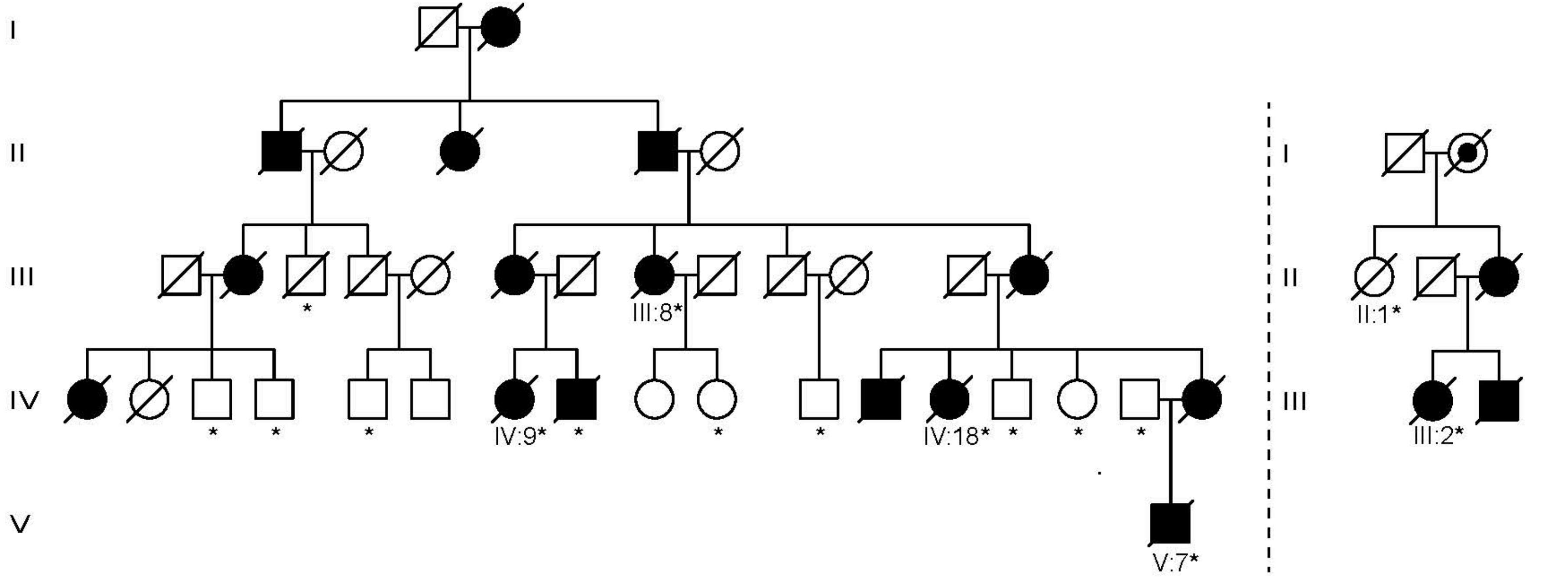


**Supplementary Figure 3. Pedigrees of families FALS5 and FALS14.** Pedigrees of family FALS5 (left) and FALS14 (right) showing segregation of the *UBQLN2* c.1460C>T (p.T487I) mutation. DNA from individuals III:8, IV:9, and IV:18 from FALS5; and individuals II:1 and III:2 from FALS14 were analysed for identity-by-descent. Brain tissue from individuals IV:18 and V:7 from FALS5 was analysed for neuropathology. Filled symbols, affected by ALS and/or FTD; Open symbols, unaffected; Asterisks, genotyped for *UBQLN2* T487I. Sex and affected status of individuals in recent generations has been masked to preserve confidentiality. Original pedigrees were published in [1].

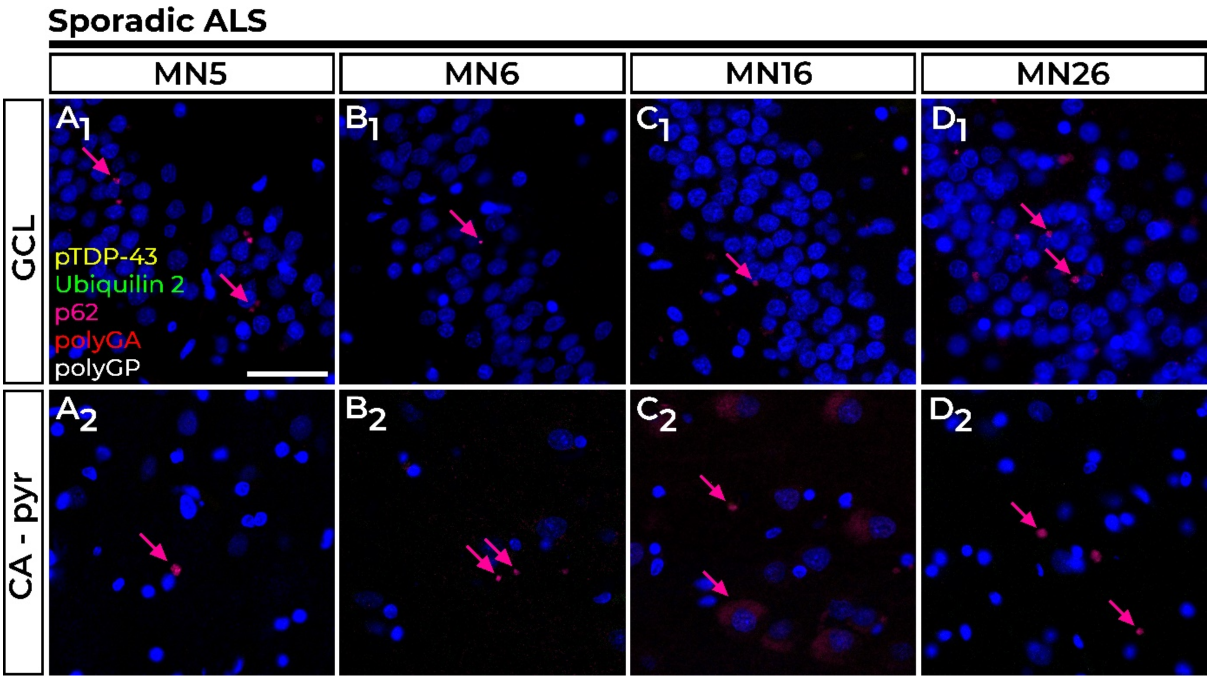


**Supplementary Figure 4. P62 pathology in the hippocampal dentate gyrus and cornu ammonis regions of sporadic ALS cases.** Punctate p62-positive inclusions were found in the GCL (**A1-D1**), and CA-pyr layers (**A2-D2**) in MN5, MN6, MN16, and MN26, all ALS cases of unknown genotypic cause. No hippocampal ubiquilin 2, pTDP-43, or DPR (polyGA and polyGP) protein pathology was observed in any of these cases. Scale bar, 50 µm.

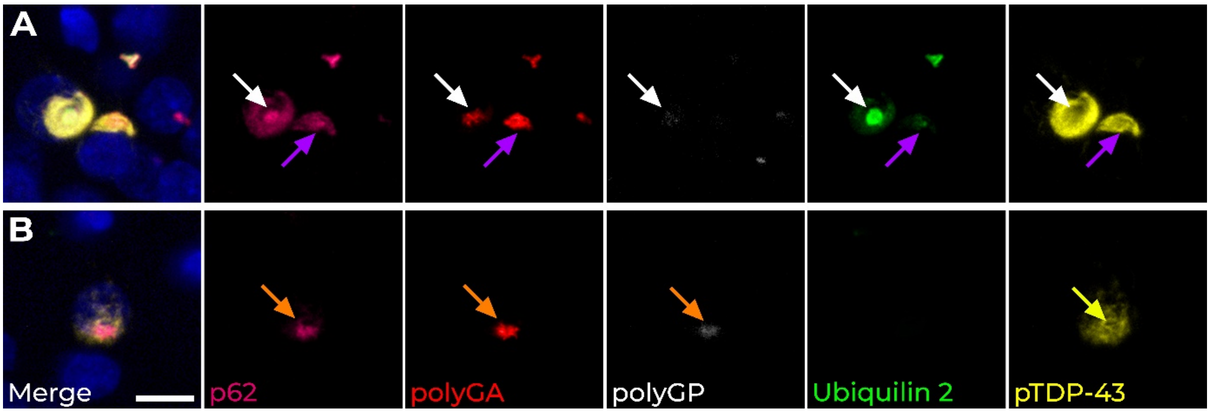


**Supplementary Figure 5. Hippocampal pTDP-43 rarely co-localises with p62-positive ubiquilin 2 and polyGA and/or polyGP in *C9orf72*-linked case MN28.** In the hippocampal granule cell layer of *C9orf72*-linked case MN28, two types aggregates in the GCL of *C9orf72*-linked case MN28 were observed to be positive for pTDP-43; one was immunopositive for all markers (**A**, white arrows), while the other was immunopositive for p62, polyGA, and ubiquilin 2, and pTDP-43 without polyGP (**A**, purple arrows). Another rare form of dipeptide repeat aggregate was immunopositive for p62, polyGA and polyGP (**B**, orange arrows) with a pTDP-43 shell (**B**, yellow arrow). Scale bar, 10 µm.

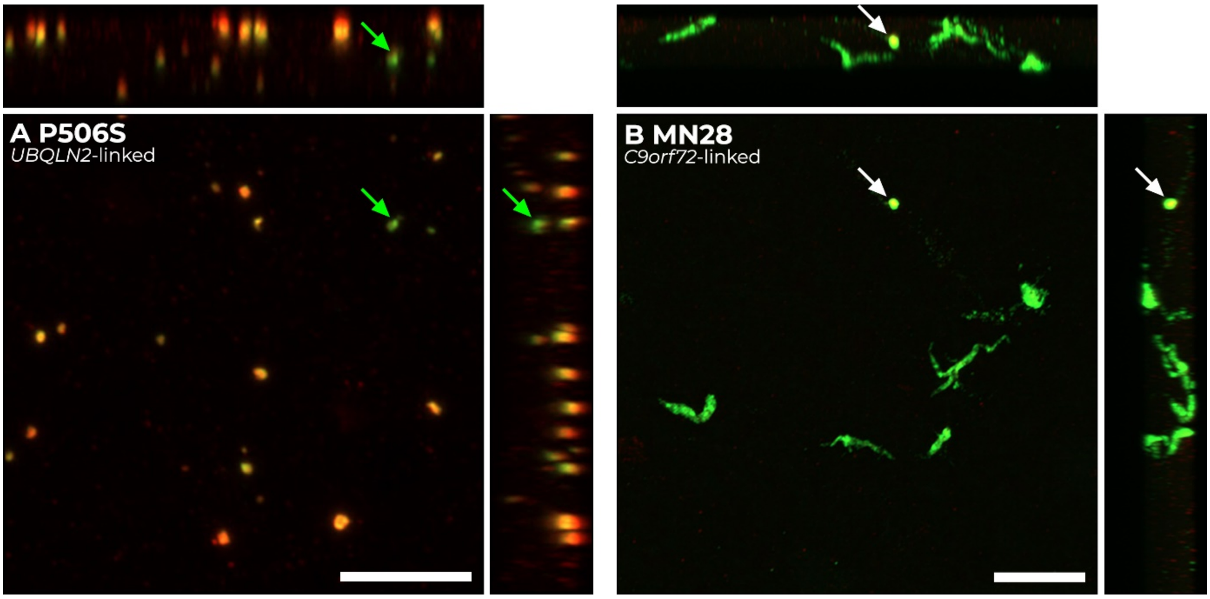


**Supplementary Figure 6. Mutant ubiquilin 2 was p62 positive in hippocampal molecular layer.** Maximum intensity Z-projections with orthogonal planes of the molecular layer confirmed that almost all mutant ubiquilin 2 aggregates in *UBQLN2*-linked ALS/FTD case p.P506S were compact and p62-labelled (**A**, z= 22.25 µm) with very few ubiquilin 2 aggregates that were p62-negative (**A**, green arrow), while the majority of wildtype ubiquilin 2 in *C9orf72*-linked case MN28 were wispy and p62-negative (**B**, z= 13 µm). A rare p62 co-labelled ubiquilin 2 aggregate is indicated with a white arrow in **B**. Scale bar, 10 µm.

**Supplementary Table 4. FALS 5 and FALS 14 relatedness analysis sample information**

| Family ID | Pedigree ID | Sex | Disease Status | *UBQLN2* p.T487I |
|---|---|---|---|---|
| FALS5 | III:8 | Female | Affected | Yes |
| FALS5 | IV:9 | Female | Affected | Yes |
| FALS5 | IV:18 | Female | Affected | Yes |
| FALS14 | II:1 | Female | Unaffected | No |
| FALS14 | III:2 | Female | Affected | Yes |
